# Pericytes Directly Communicate with Emerging Endothelial Cells During Vasculogenesis

**DOI:** 10.1101/2020.07.01.180752

**Authors:** Laura Beth Payne, Bhanu Tewari, Logan Dunkenberger, Samantha Bond, Alyssa Savelli, Jordan Darden, Huaning Zhao, Michael Powell, Kenneth Oestreich, Harald Sontheimer, Sophie Dal-Pra, John C. Chappell

**Author notes:** Previous affiliation(s) (Logan Dunkenberger, Alyssa Savelli, Jordan Darden, Huaning Zhao, Sophie Dal-Pra). Corresponding Author: John C. Chappell, Ph.D. Fralin Biomedical Research Institute at Virginia Tech-Carilion 2 Riverside Circle Roanoke, Virginia 24016 E-mail Correspondence.

## Abstract

Pericytes (PCs), cells that extend along capillaries to contribute stability and other critical functions to established vasculature, are attracting attention from various fields involving vascular-related pathologies. Here, we demonstrate primary evidence of PC communication with endothelial cells (ECs) prior to tube coalescence. Observations of apparent PCs during early embryogenesis urged development of a mouse embryonic stem cell line (DR-ESCs), enabling unique dual-reporter investigations into earliest PC-EC interactions. Live imaging of differentiating DR-ESCs corroborated emergence of a PC lineage, which preceded EC differentiation, and further revealed highly dynamic PC-EC interactions during coordinated vessel formation. We show direct PC-EC communication via cell microinjection and dye-transfer, and RNA-seq analysis indicates a PC-EC coupling mechanism via gap junction Connexin43 (Cx43), exclusively up-regulated throughout DR-ESC differentiation. High resolution imaging of embryonic and postnatal mouse vasculature substantiates Cx43 plaques at PC-EC borders. These findings indicate a new role for PCs during vasculogenesis via Cx43-mediated communication with ECs.

## INTRODUCTION

Pericytes (PCs) extend along and stabilize capillaries within microvascular networks of many organs and tissues. Vascular PCs establish non-overlapping domains on vessels throughout the microcirculation (Grant et al., 2017) and have several established and proposed roles, such as blood flow regulation (Hall et al., 2014; Hill et al., 2015), shaping the extracellular matrix (ECM) surrounding blood vessels (Fernandez-Klett et al., 2013; Sava et al., 2015), and enhancing endothelial cell (EC) junction formation (Jo et al., 2013), in normal and pathological environments. These functions promote and maintain vascular integrity of established vasculature and thus, regulate vessel barrier function (von Tell et al., 2006; Winkler et al., 2018; Zhao et al., 2018). PCs directly interact with established endothelium via “peg-and-socket” adhesion molecules (Diaz-Flores et al., 2009) and intercellular communication channels of connexin-based gap junctions (GJs) (Fang et al., 2013; Hirschi et al., 2003). While, PC-EC point contacts have been observed in cell culture models and within mature vessels, including sprouting angiogenesis, it is unclear how early in vascular development PCs interact with ECs.

It is widely thought that PC differentiation and vessel association occurs later in development, perhaps to allow for greater EC remodeling plasticity (Bergers and Song, 2005; Marmé and Fusenig, 2008). It has been suggested that PCs arise from the EC compartment (Chen et al., 2016) and may be dependent on EC interactions (Fang et al., 2013; Hirschi et al., 2003), implying that PCs differentiate after the endothelium is largely established. However, PCs have been largely unexamined during early angiogenesis or vasculogenesis. In part, this stems from challenges inherent in *in vivo* investigations at these early time points, and a limit of complementary models that enable investigation of PC differentiation and PC-EC interactions in vasculogenesis. However, some recent advances have been made.

Embryonic (ESCs) and induced-pluripotent stem cells (iPSC) have proven to be relevant models for illuminating fundamental developmental and functional processes of various tissues systems, including vascular. Development of mesoderm-derived hematopoietic, vascular, and cardiac lineages from differentiating ESCs occurs robustly in serum-exposed EBs (Keller, 2005), giving rise to mural cells (D’Souza et al., 2005; Ema et al., 2003), and ECs that reflect initial *in vivo* differentiation in yolk sac blood islands (Haar and Ackerman, 1971) (Vittet et al., 1996). Although ESCs lack blood flow, ESC-derived ECs form organized yet primitive vessels that lumenize and provide insight into vessel wall and network assemblies that serve as templates for further remodeling (Bautch et al., 1996; Doetschman et al., 1985; Lucitti et al., 2007; Yamashita et al., 2000). While PC origin (Faal et al., 2019) (Trost et al., 2016) (Volz et al., 2015) (Chen et al., 2016) and growth factors that influence differentiation (Armulik et al., 2011) have been described, the mechanisms underlying emergence are still being established. We previously used mouse ESCs to explore PC-EC decoupling downstream of aberrant VEGF-A signaling during sprouting angiogenesis (Darden et al., 2018). Here, we present a validated, double-reporter mouse ESC line that contains reporters for ECs (*Flk-1:eGFP*) and PCs (*Ng2/Cspg4:DsRed*), as a new tool in which to investigate and quantify real-time dynamics of early stage vessel development.

Although PCs have gained considerable interest recently, there remain crucial gaps in knowledge regarding PC function, including differential spatial and temporal roles. Using DR-ESCs to complement *in vivo* mouse studies, we examined PC-EC interactions and differentiation dynamics at early-stage vascular development, testing the hypothesis that PCs arise and engage with ECs during vasculogenesis. We found that *Ng2-DsRed*+ PCs arise prior to mesoderm-derived *Flk-1:eGFP*+ ECs and tube formation, from distinct precursors. Moreover, we demonstrate direct PC communication with ECs during vasculogenesis, and put forth evidence for Connexin43 (Cx43)-mediate gap junction mechanism of cell-cell transfer. Our findings here support a new role for PCs in coordinated vessel formation, and provide new insights into the development of functional vasculature.

## RESULTS

### Pericytes are Apparent During Vasculogenesis in Mouse Embryos

Using confocal microscopy, we examined fresh, whole mouse embryos that harbored fluorescent reporters for ECs (*Flk-1-eGFP*) and PCs (*NG2-DsRed*) at embryonic day 8.5 (E8.5), a developmental stage when ECs were present but not yet coalescing into vessels (Figure 1A). We found distinct *Ng2:DsRed*+ cells (Figure 1A ii, v, viii) among *Flk-1:eGFP+* ECs that appeared to be arising from *Flk-1:eGFP+* mesoderm (Figure 1A, i, iv, vii). At E9.5 we found *Ng2-DsRed*+ cells along the heart outflow tract and adjacent regions of the dorsal aorta – structures that form in part via vasculogenic processes (Figure 1B and <u>Online Video S1</u>). Although oligodendrocyte progenitor cells also express NG2, this occurs later in development (Trotter et al., 2010), indicating that the *Ng2:DsRed*+ cells observed in these E8.5 and E.9.5 embryos are PCs or related mural cells. This was surprising, as PCs are thought to arise later in development and engage with ECs following vessel structure formation (Bergers and Song, 2005; Marmé and Fusenig, 2008). Therefore, in order to broadly probe these intriguing findings, we developed an embryonic stem cell model to complement *in vivo* studies.

**Figure 1.**
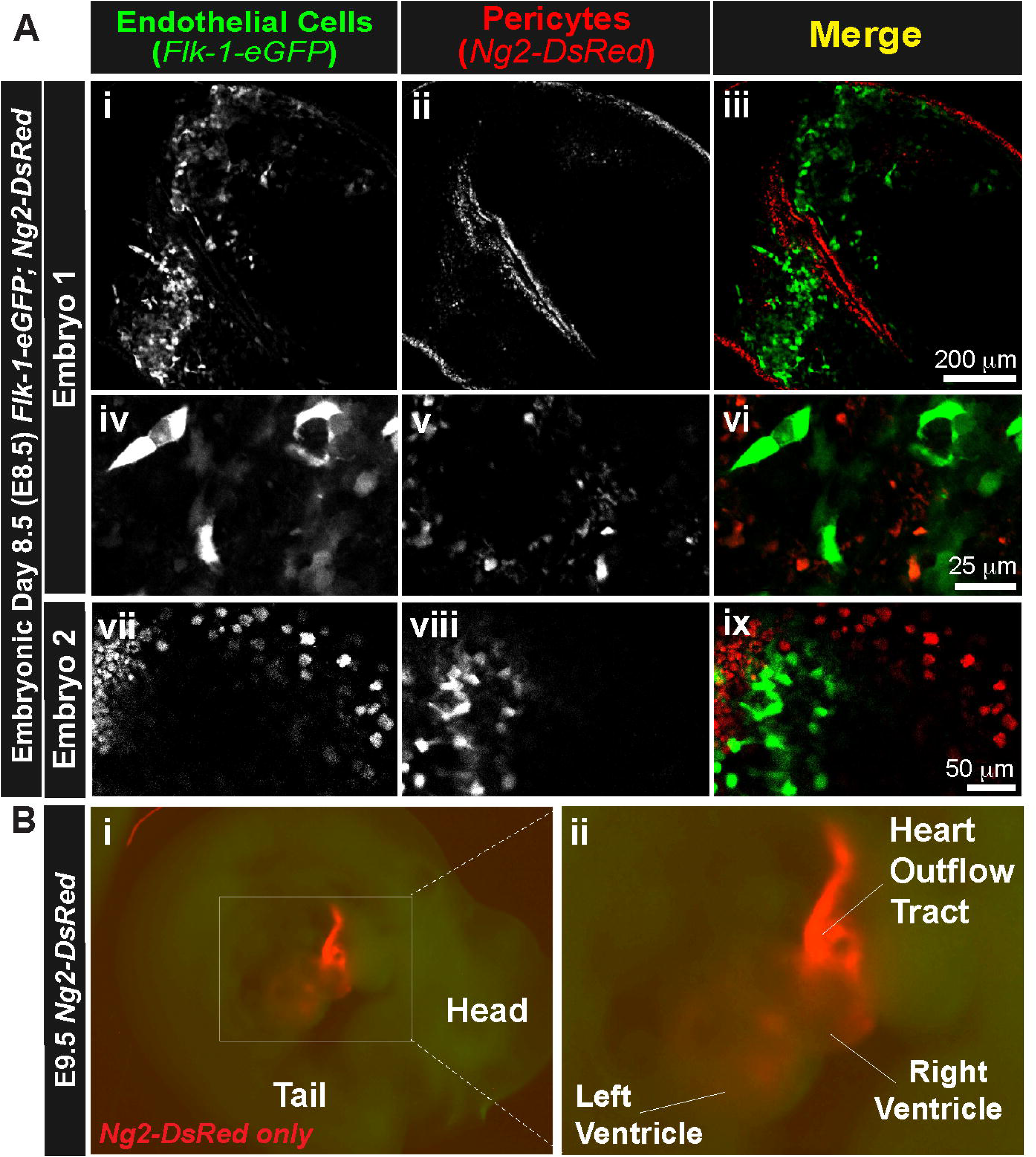
Pericytes are Apparent During Vasculogenesis in Mouse Embryos. **(A)** Single plane confocal images of embryonic day 8.5 (E8.5) whole mouse embryos (embryo 1 – i-iii and iv-vi, embryo 2 – vii-ix), with endogenous reporters for ECs, *Flk-1-eGFP* (green), and PCs, *Ng2-DsRed* (red) **(B)**: Heart outflow tract of an E9.5 *Ng2-DsRed+* embryo. (i) Epi-fluorescent microscope image of an embryonic day 9.5 (E9.5) heart from a *Flk-1-eGFP-negative; Ng2-DsRed-positive* mouse embryo, demonstrating mural cells along the heart outflow tract (red) with gross animal morphology (dim green); (ii): Higher magnification of the *Ng2-DsRed+* mural cells along the heart outflow tract. *See **<u>Online Video S1</u>***.

### A Genetically Stable and Highly Proliferative EC- and PC-Reporter ESC Line Maintains Pluripotency, Differentiating into Each Germ Layer

To establish DR-ESCs as a viable model, we verified critical characteristics (Gaztelumendi and Nogues, 2014). We collected blastocysts at E3.5 from a mouse with reporters for ECs and PCs (*Flk-1:eGFP;NG2-DsRed*). We maintained pluripotency in LIF, 10% serum and dual inhibition of signaling kinases mitogen-activated protein kinase kinase (MEK) and glycogen synthase kinase-3 (GSK3) (i.e. the “2i system”) – an approach known to enhance uniform pluripotency (Wray et al., 2010). We confirmed genetic stability and absence of chromosomal abnormalities, such as aneuploidy, via karyotyping (Figure S1A). Undifferentiated DR-ESCs were uniformly positive for alkaline phosphatase staining (Figure S1B), verifying pluripotency (O’Connor et al., 2008). To initiate differentiation, DR-ESCs aggregated into embryoid bodies (EBs), followed by removal of LIF and 2i inhibition. By days 5 and 12 of differentiation, alkaline phosphatase staining progressively diminished into small cell clusters throughout the differentiating EBs (Figure S1B). Next, we confirmed DR-ESC proliferative capacity via immunostaining for phospho-histone H3 (PH3), a mitotic indicator. Undifferentiated and day 5 differentiated cultures contained high numbers of PH3+ cells (Figure S1C), indicating their potential for expansion via robust cell division.

We evaluated pluripotent stem cell markers via immunocytochemistry (ICC) and gene expression. Positive Oct4 immunostaining (Czechanski et al., 2014) in undifferentiated cells was wide-spread (Figure S1D). After 5 and 12 days of differentiation, the number of Oct4+ cells progressively decreased and localized into discrete pockets (Figure S1D). We confirmed down-regulation of transcript expression of stem cell markers, *Oct4, Fgf4, Sox2*, *Nanog* and *c-kit/CD117,* via qRT-PCR in days 5 and 12 differentiated DR-ESCs (Figure S1E). Gradual loss of stem cell signatures coincided with the rise of each germ layer – mesoderm, ectoderm and endoderm. We analyzed transcript levels of germ layer markers *Dab2, Gata6, Actc1* and *Otx2*. Expression of stem cell markers was significantly higher in the undifferentiated DR-ESCs as compared to days 5 and 12 of differentiation (Figure S1E), while germ layer marker expression dramatically increased from undifferentiated to day 5 of differentiation, with mesodermal markers showing high expression (Figure S1F). Germ layer markers increased further at day 12, with the exception of a decrease in the ectoderm marker *Otx2*. RNA sequencing (RNA-Seq) of additional germ layer markers in undifferentiated, days 7 and 10 differentiated ESCs supported these quantifications (Figure S2).

### DR-ESC Differentiation Yields Vascular EC and PC Lineages

We verified the specificity of the reporter genes, specifically *Flk-1:eGFP* in ECs and *Ng2:DsRed* in PCs. To do this, we examined transcript expression in undifferentiated, days 5 and 12 differentiated DR-ESCs for EC markers: *Pecam1/CD31, VE-Cadherin/Cdh5/CD44, Flk-1/Kdr/Vegfr2, Icam2/CD102*, and for accepted PC markers (Armulik et al., 2011): *Ng2/Cspg4, Pdgfrb*/*CD140b*, *Desmin*, and *N-Cadherin/Cdh2* (Figure 2A & B). With the exception of *Pecam1/CD31*, each gene steadily increased from little to no expression in undifferentiated cells through days 5 and 12 differentiation (Figure 2A & B). Expression of *Pecam1/CD31* in undifferentiated mouse ESCs aligns with previous studies (Li et al., 2005; Robson et al., 2001). Also significantly up-regulated over the 12 days of differentiation was *Tgfb1* (Figure 2A), a mesodermal germ layer marker and a key stimulus for inducing PC and vascular smooth muscle cell (vSMC) differentiation (Gaengel et al., 2009; Hirschi et al., 2003; Sieczkiewicz and Herman, 2003). Overall, these transcription profiles demonstrate robust EC and PC differentiation within DR-ESCs over time.

**Figure 2.**
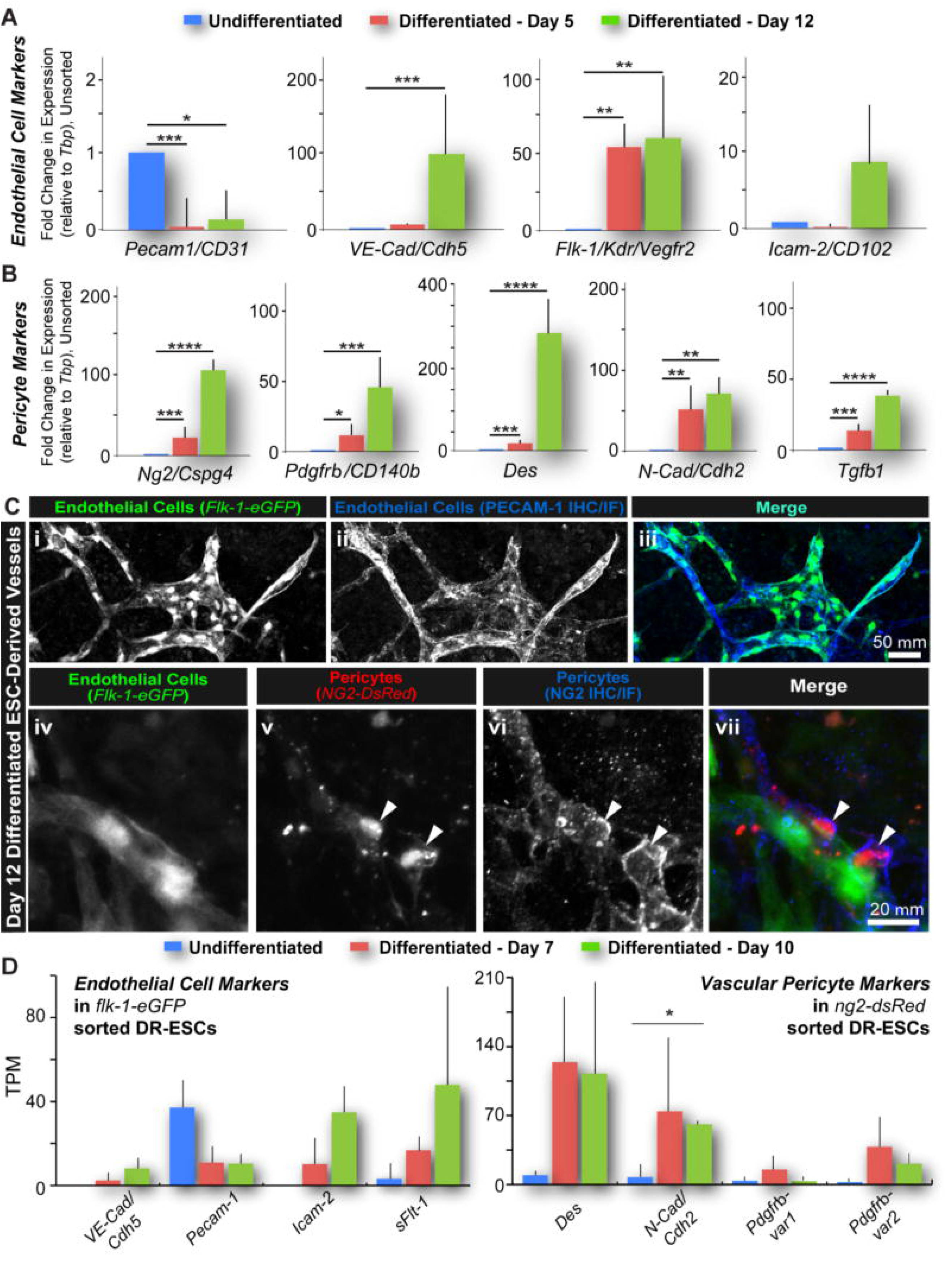
DR-ESC Differentiation Yields Vascular EC and PC Lineages. **(A):** EC marker expression from unsorted undifferentiated, days 5 and 12 differentiated DR-ESCs. **B):** Relative PC marker expression and T*gfβ1* (PC differentiation cue) from unsorted undifferentiated, days 5 and 12 differentiated DR-ESCs. *P≤0.05, **P≤0.01, ***P≤0.001, ****P≤0.0001. Error bars, SEM (n=3). **(C):** Day 12 differentiated DR-ESC-derived ECs (*Flk-1-eGFP*+; i; iii) labeled for PECAM-1 (ii, iii). NG2-DsRed+ PCs (v; vii) along *Flk-1-eGFP*+ ECs (iv; vii) labeled for cell surface NG2 protein (vi; vii). **(D):** Cell- specific expression of vascular markers by RNA-Seq for FACS-sorted ECs *(Flk-1-eGFP+*) and PCs (*Ng2-DsRed+*) in undifferentiated, days 7 and 10 differentiated DR-ESCs. TPM, Transcripts per million. *P≤0.05. Error bars, SD (n=2). *See* ***Figure S3*.**

We verified that the reporters labeled the expected cell populations via ICC and confocal imaging. We immunostained day 12 differentiated DR-ESCs for PECAM-1 and NG2 protein (Figure 2C). The PECAM-1 signal consistently overlapped with eGFP, as expressed by the *Flk-1* promoter, confirming that eGFP+ cells also expressed this EC marker (Figure 2Ci-iii). Cells expressing *Ng2/Cspg4*, as indicated by DsRed fluorescence, were also positively labeled for cell surface NG2 (Figure 2Civ-vii). We confirmed transcriptional profiles of cell-specific gene expression over time via RNA-seq analysis. We sorted *Ng2:DsRed*+ and *Flk-1:eGFP*+ populations using fluorescence-activated cell sorting (FACS). Prior to FACS, we used flow cytometry to confirm the lack of reporters in undifferentiated DR-ESCs (Figure S3A), as well as non-overlap of signals throughout differentiation (Figure S3A). We visually confirmed the expected reporter signals before and after FACS, using epi-fluorescent microscopy (Figure S3B). Sorted PC and EC populations from undifferentiated, days 7 and 10 differentiated DR-ESCs were processed via paired-end RNA-seq (Genewiz, NJ) and alignment to the *mus musculus* genome assembly, GRCm38.p6. Transcript expression, analyzed (Kallisto (Bray et al., 2016)) for *Flk-1:eGFP*+ ECs and *Ng2:DsRed*+ PCs (Figure 2D), were consistent with expected markers and with the candidate approach applied to unsorted cells (Figure 2A & B).

Due to the lack of a singular PC marker (Armulik et al., 2011), they are most confidently identified through a combination of labels and morphological features. We used ICC to co-label DsRed+ cells in day 6-8 differentiating cultures for PC markers PDGFRb (Figure 3Ai-iv) and desmin (Figure 3Av-viii). We also excluded the possibility of DsRed fluorescence being retained within daughter cells, by confirming active NG2 protein synthesis at days 6 (Figure 3Aix-xii) and 12 (Figure 2Civ-vii). We distinguished vessel-associated *Ng2:DsRed*+ cells from NG2-expressing oligodendrocyte progenitor cells (OPCs) (Polito and Reynolds, 2005) via ICC with A2B5 (Figure 3B). A2B5 signals were detected: (i) nearby, but not coinciding with, *Flk-1:eGFP+* vessels (Figure 3Bv-xii), and (ii) with no apparent co-labeling of *Ng2:DsRed*+ cells. We examined transcript levels of oligodendrocyte markers *Mbp*, *Olig1, Olig2,* and *Plp1* from RNA-seq data of *Ng2:DsRed+* sorted DR-ESCs. At days 7 and 10 of differentiation, levels of these markers were below a 5 transcripts per million (TPM) cutoff (data not shown), further distinguishing *Ng2:DsRed+* cells from glial lineages.

**Figure 3.**
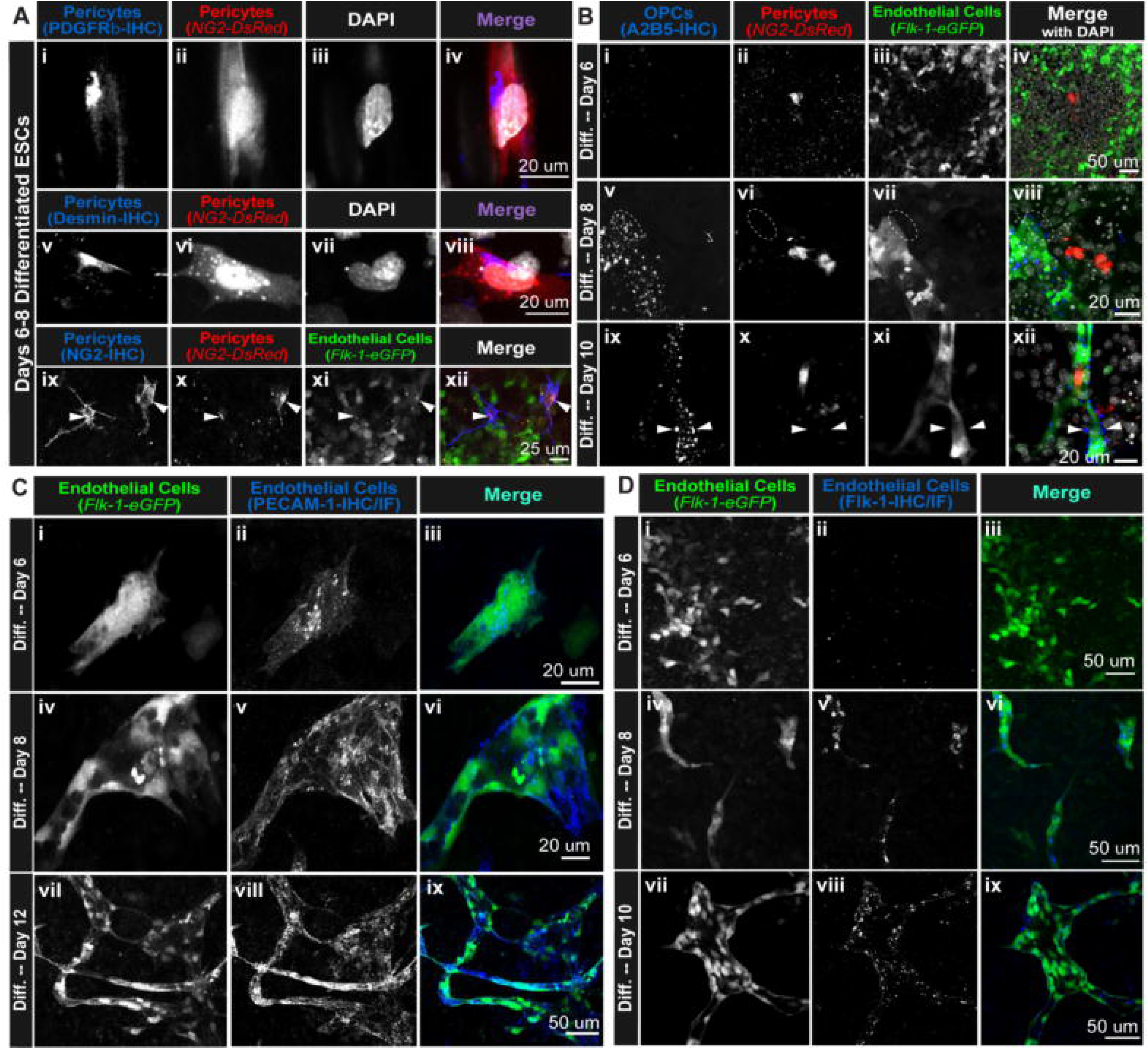
Reporter Constructs Coincide with Vascular EC and PC Markers throughout DR-ESC Differentiation. **(A):** Days 6-8 differentiated DR-ESC-derived PCs (*Ng2-DsRed+*; ii, vi, x; iv, viii and xii) labeled for PDGFRβ (i; iv), Desmin (v; viii), and NG2 (ix; xii). *Flk-1-eGFP+* ECs (xi; xii). Arrowheads indicate co-labeled cells (ix-xii). **(B):** DR-ESC-derived PCs (*Ng2-DsRed+*; ii, vi, x; iv, viii and xii) not labeled by the OPC marker A2B5 (i, v, ix; iv, viii and xii) in days 6 (i-iv), 8 (v-viii), 10 (ix-xii) differentiated DR-ESCs. *Flk-1-eGFP+* mesoderm (iii) and ECs (vii, xi; iv, viii, xii). Dotted ovals (v-viii) and arrowheads (ix-xii) note A2B5 signal adjacent to, but not overlapping with, *Flk-1-eGFP*+ ECs. **(C):** DR-ESC-derived ECs (*Flk-1-eGFP+*; i, iv, vii; iii, vi and ix) labeled for PECAM-1 (ii, v, viii; iii, vi, and ix) from days 6 (i-iii), 8 (iv-vi), 12 (vii-ix) of differentiation. **(D):** DR-ESC-derived ECs (*Flk-1-eGFP+*; i, iv, vii; iii, vi ix) labeled for Flk-1 receptor from days 6 (i-iii), 8 (iv-vi), 10 (vii-ix) of differentiation. S*ee* ***Figure S4*.**

We profiled DR-ESC-derived *Flk-1:eGFP*+ cells using ICC to label PECAM-1 in early-, mid-, and late-stage endothelial differentiation (days 6, 8 and 12) to corroborate EC identity throughout differentiation (Figure 3C) and to distinguish ECs from precursors. We labeled Flk-1 on the EC surface over time (days 6, 8 and 10) to verify active Flk-1 production from *Flk-1:eGFP+* cells (Figure 3D). Both PECAM-1 and Flk-1 staining co-localized with highly-expressing *Flk-1:eGFP* cells (denoted *Flk-1-eGFP^high^*) at each time point, delineating eGFP^high^ cells as ECs. We corroborated these observations via spatial line-scan analysis of PC, OPC, and EC markers with respect to *NG2-DsRed*+ and *Flk-1-eGFP*+ fluorescence (Figure S4).

### DR-ESC-Derived ECs and PCs Organize into Primitive Vascular Networks in Proximity to Contracting Cardiomyocytes

Using short-term, time-lapse imaging, we identified nascent cardiac tissue by observing synchronous contraction of early-stage cardiomyocytes (<u>Online Video S2</u>). Overlaying the fluorescent signal from the *Flk-1:eGFP+* ECs and *Ng2:DsRed+* PCs, we observed early vessels in close proximity to primitive cardiac tissue (Figure 4A), recapitulating another key feature of early vascular development. We found that many regions of developing cardiomyocytes contained or were neighboring DR-ESC-derived vessels. Vessels were also apparent at distal and spatially heterogeneous locations (Figure 4A), suggestive of primitive capillary networks that arise in other embryonic and extra-embryonic tissues during development.

**Figure 4.**
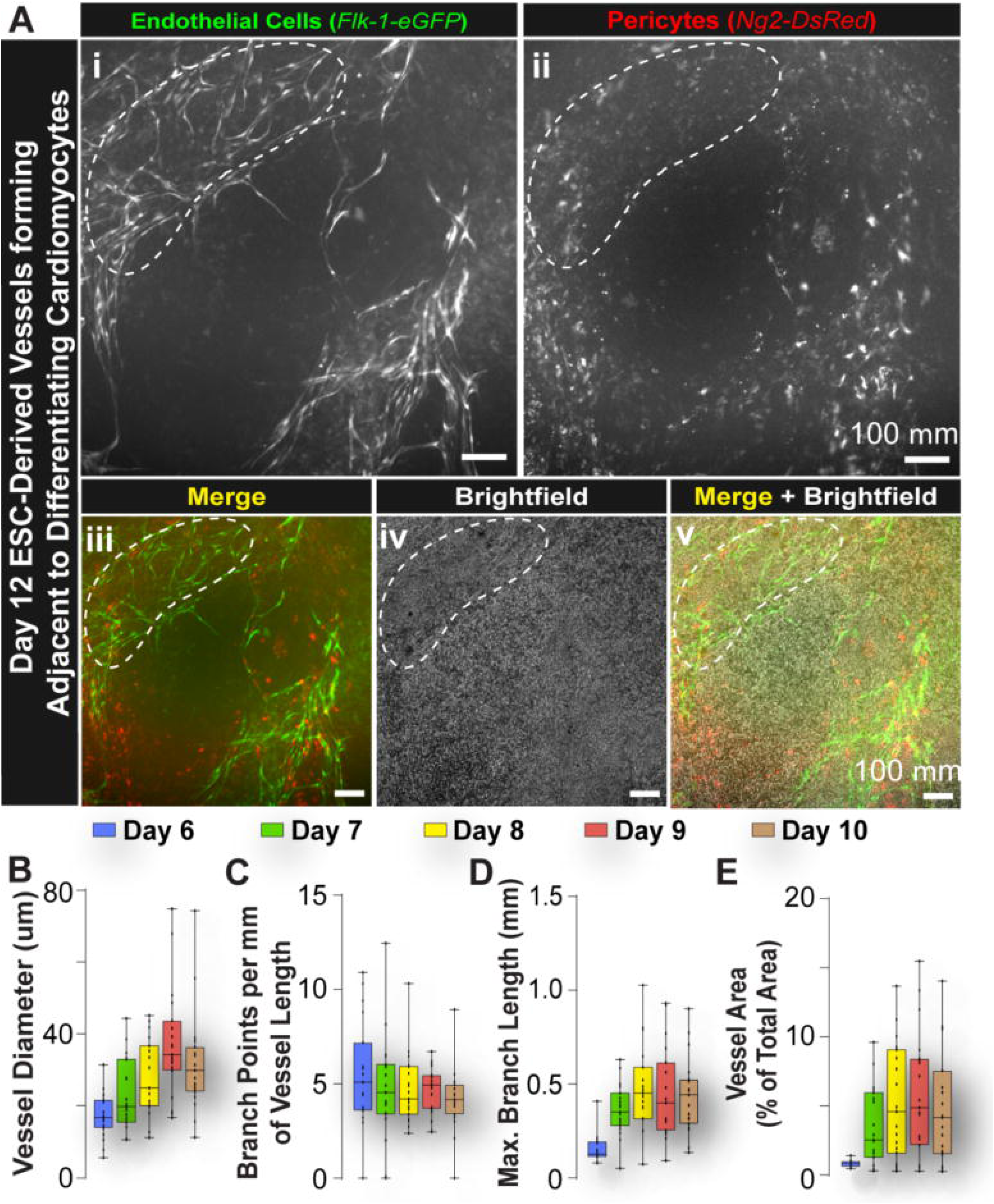
DR-ESC-Derived ECs and PCs Organize into Primitive Vascular Networks in Proximity to Contracting Cardiomyocytes. **(A):** *Flk-1-eGFP+* ECs (i; iii, v) and *Ng2-DsRed*+ PCs (ii; iii, v) in vessels near contracting cardiomyocytes (white dashed line) and neighboring areas. *See **<u>Online Video S2</u>***. **(B-E):** Morphological quantifications of DR-ESC-derived vessels at days 6-10 of differentiation - vessel diameter **(B)**, branch points per mm of vessel length **(C)**, branch length (mm) **(D)**, and vessel area as a percent of total area **(E)**.

We quantified dynamic formation of vessel networks via long-term, dynamic confocal imaging from day 6 to day 11 of differentiation, using Image J (FIJI) software (Schindelin et al., 2012). Analysis revealed increases in: 1) vessel diameter (Figure 4B), 2) maximum vessel branch length (Figure 4D), and 2) projected vessel area (as a percentage of the total area within a field-of-view) (Figure 4E) followed by a plateau or slight regression. Branch points per mm of vessel (Figure 4C) remained steady over time, indicating biphasic growth via initial expansion and subsequent stabilization.

### PCs Differentiate from Distinct Precursors Prior to ECs

Using high resolution confocal microscopy, we observed a notable increase in eGFP fluorescence (*Flk-1:eGFP^high^*) during EC differentiation from precursor mesoderm that exhibited a distinctly lower *Flk-1:eGFP* signal (*Flk-1:eGFP^low^*; <u>Online Video S2</u>). To confirm delineation of these two populations, we labeled PECAM-1 via ICC within early-stage ESC differentiations, where high and low eGFP signals were simultaneously present (Figure 5A). The PECAM-1 staining exclusively labeled *Flk-1:eGFP^high^* cells, distinguishing these differentiating ECs from the surrounding mesoderm (*Flk-1:eGFP^low^*) from which they arise (described below).

**Figure 5.**
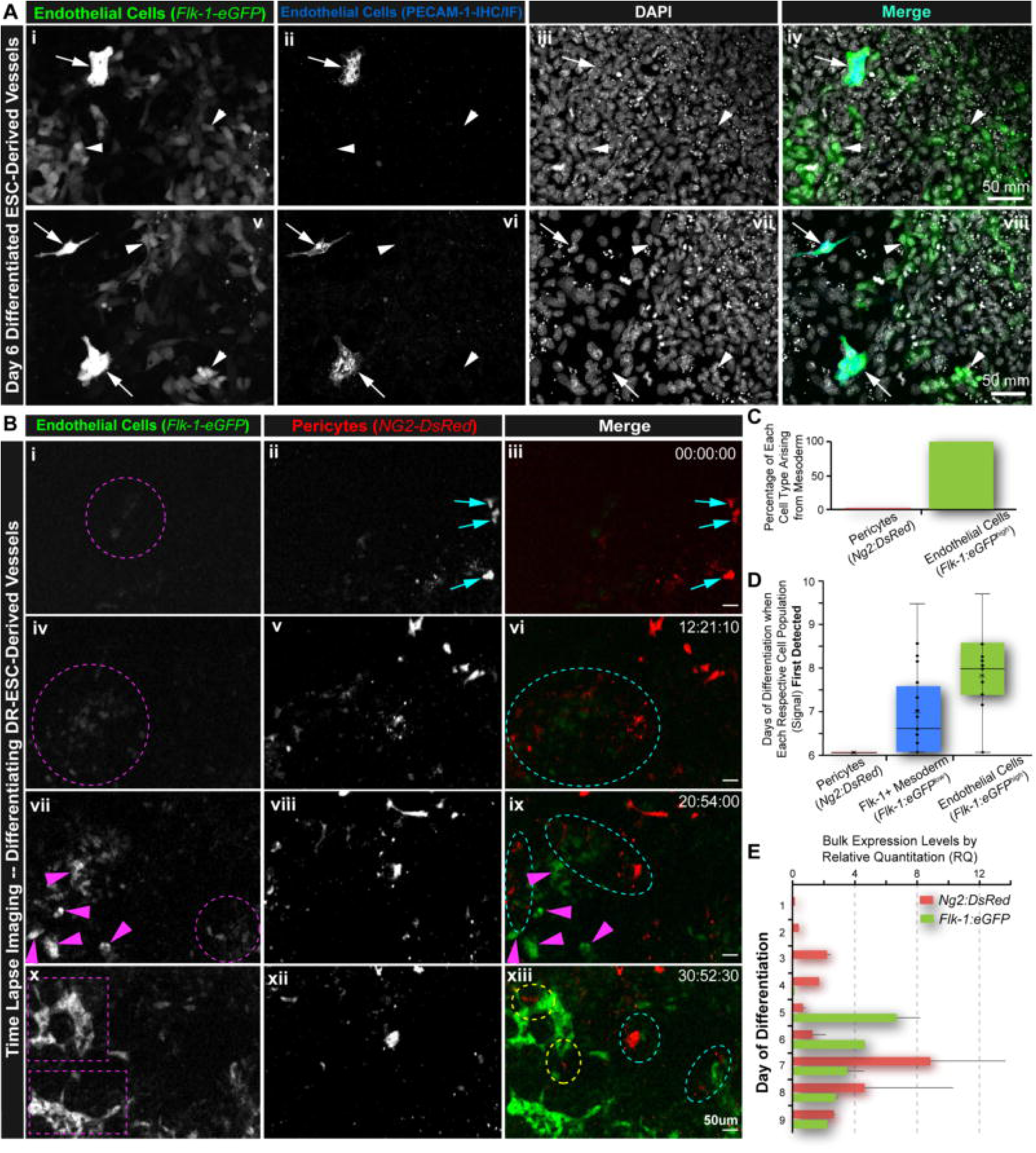
PCs Differentiate from Distinct Precursors Prior to ECs. **(A):** Day 6 differentiated DR-ESCs (i-iv, v-viii) labeled for PECAM-1 (ii, vi; iv, viii). Arrows note *Flk-1-eGFP^high^* ECs (i, v; green^high^ – iv, viii). Arrowheads note precursor *Flk-1-eGFP^low^* mesoderm (i, v; green^low^ – iv, viii). Nuclei, DAPI (iii, vii; iv, viii). See ***<u>Online Video S3</u>***. **(B):** Time-lapse images of DR-ESC differentiation over ∼30 hours. Magenta arrowheads (vii, ix) denote emergence of ECs (x; xii). Dashed magenta circles are precursor *Flk-1-eGFP^low^* mesoderm (i, iv, vii; iii, vi, ix). Dashed purple boxes indicated EC coalescence into vessels (x). Cyan arrows are *Ng2-DsRed+* PCs (ii, v, viii, xi; iii, vi, ix, xii) arising before *Flk-1-eGFP+* signals. Dashed cyan ovals note *Ng2-DsRed+* PC and *Flk-1-eGFP+* mesoderm interactions (vi, ix, xii). Dashed yellow ovals note *Ng2-DsRed+* PC participating in EC organization (xii,). Time (hh:mm:sec). *See **<u>Online Video S4</u>**.* **(C):** Initial detection of Ng2-DsRed+ PCs, Flk-1-eGFP+ mesodermal precursor cells (*Flk-1-eGFP^low^*), and *Flk-1-eGFP+* ECs (*Flk-1-eGFP^high^*) in days 6 to 11 differentiating DR-ESC (n=21, blinded). **(D):** Percentages of *NG2-DsRed+* PCs and *Flk-1-eGFP^high^* ECs arising from *Flk-1-eGFP^low^* mesoderm (n=21, blinded). **(E):** Relative quantitation (RQ) of *DsRed* (*Ng2*) and *eGFP* (*Flk-1)* transcripts from unsorted DR-ESCs at day 1 (undifferentiated) through 9 of differentiation. Error bars, SD, (n=2).

#### Non-overlapping progenitor populations for PCs and ECs

To examine whether PCs and ECs may arise from a common *Flk-1:eGFP+* progenitor, (Chen et al., 2016; Yamashita et al., 2000), we assessed time-lapse imaging of early stage DR-ESC differentiation and did not observe *Flk-1:eGFP+* precursor cells generating double-positive PCs (i.e. *Flk-1:eGFP+* and *Ng2:DsRed+* cells) (Figure 5B). We quantified percentages of both DsRed+ cells and eGFP^high^ ECs that arose from *Flk-1:eGFP+* precursor cells (Figure 5C) in blind analysis. All observed *Flk-1:eGFP^high^* ECs were derived from *Flk-1:eGFP+* mesodermal precursors. In contrast, we did not detect a single occurrence of DsRed+ fluorescence overlapping with a *Flk-1:eGFP+* cell, suggesting that PCs and ECs in our model did not arise from a common Flk-1+ precursor.

#### Temporal differentiation of PCs and ECs

We quantified time-lapse imaging of DR-ESC differentiation from day 6 through 10, aiming to detect the earliest phases of vasculogenesis and the emergence of each cell type. Our analysis revealed distinct signals from the *Ng2:DsRed* reporter at the earliest time point captured during live imaging (day 6), which consistently appeared prior to *Flk-1:eGFP^high^* signals (Figure 5B and <u>Online Video S3</u>). We quantified mRNA transcripts of *DsRed* and *eGFP* via qRT-PCR in undifferentiated cultures (day 1) through day 9 differentiation (Figure 5E). Increased *DsRed* transcript levels occurred on day 3 as compared to *eGFP* transcripts, which first appeared on day 5 and presumably represents abundant *Flk-1:eGFP^low^* mesoderm, (Figure 5B, i, iv)). These data indicate that PCs, or their precursors, arise prior to EC differentiation in this setting.

### During Vasculogenic and Angiogenic Remodeling, PCs and ECs Interact Directly via Gap Junctions Prior to Tube Formation

#### Direct PC-EC Interactions during Vasculogenesis

Consequent to *Ng2:DsRed+* PC emergence, we acquired long-term movies of early stage differentiation encompassing vasculogenesis (days 5-8), and found *Ng2:DsRed*+ PCs (and/or their precursors) that appeared to directly and dynamically interact with *Flk-1:eGFP*+ cells (Figure 6A, Figure S5, and <u>Online Video S4</u>, <u>Video S5</u>, and <u>Video S6</u>). We assessed the spatial proximity of neighboring PCs and ECs via orthogonal, 2.5- and 3-dimensional renderings of through-thickness confocal images at days 6 and 8 of differentiation (Figure S6), which supported direct PC-EC contact. In later-stage differentiation (i.e. days 12 and 13), we observed EC sprouting events as well as *Ng2:DsRed*+ PC migration along developing vessels (Figure 6 and <u>Online Video S7</u>), morphologically consistent with hallmarks of vascular PCs (Armulik et al., 2011). This indicates PC engagement with the developing endothelium from early-to-late stage vessel formation.

**Figure 6.**
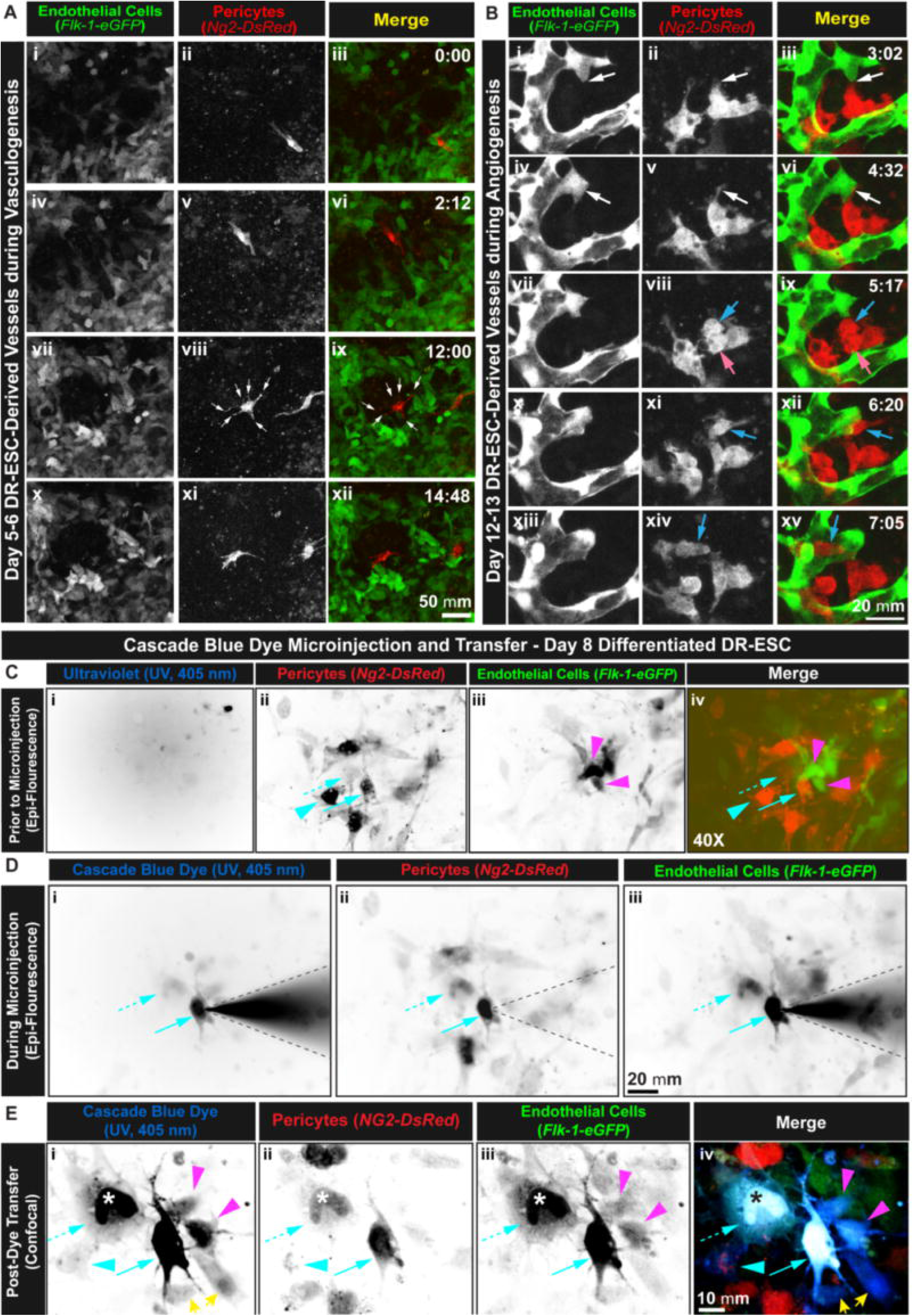
During Vasculogenic and Angiogenic Remodeling, Direct PC-EC Interaction via Gap Junctions Prior to Tube Formation. **(A)**: Time-lapse images of *Flk-1-eGFP*+ mesoderm (Flk-1-eGFP^low^ - i, iv, vii, x; iii, vi, ix, xii) and coalescing ECs (Flk-1-eGFP^high^ - vii, x, green^high^ – ix, xii) over day 5-6 of DR-ESC differentiation. *NG2-DsRed+* PCs (ii, v, viii, xi; iii, vi, ix, xii) engage with multiple ECs (white arrows – viii, ix). Time (hh:mm). *See **<u>Online Video S5</u>**,* ***Figure S5*** *and **<u>Online Video S6</u>**.* **(B):** Time-lapse images of a sprouting *Flk-1-eGFP*+ ECs (i, iv, vii, x, xiii; iii, vi, ix, and xv) interacting with *NG2-DsRed+* PCs (ii, v, viii, xi, xiv; iii, vi, ix, xii, xv; white arrows - i-vi) over days 12-13 of differentiation. Dividing PC (cyan, magenta arrows -viii, ix) yields a daughter PC (cyan arrows – xi, xii) that engages an EC sprout and migrates to nearby branch point (cyan arrows – xiv, xv). Time (h:mm). *See **<u>Online Video S7</u>** and* ***Figure S6.* (C-E)**: Cyan arrows note the 1st (solid arrow) and 2nd (dashed arrow) *NG2-DsRed+* PCs sequentially microinjected. The 2nd microinjected PC (dashed arrow) did not survive the procedure (white asterisk). Cyan and magenta arrowheads respectively note an *Ng2-DsRed+* PC and *Flk-1-eGFP+* ECs that received dye transfer. **(C):** Day 8 differentiated DR-ESCs before dye microinjection visualizing (40x) the ultraviolet (UV) spectrum (minimal signal in i, iv), *Ng2-DsRed+* PCs (ii; iv), and *Flk-1-eGFP*+ mesoderm (iii; green^low^ - iv) and ECs (iii; green^high^ - iv). **(D):** Cascade Blue dye UV emission (i), *NG2-DsRed+* PCs (ii), and *Flk-1-eGFP*+ ECs and mesoderm (iii) during the 1st PC microinjection. Dye-filled needle (dashed black lines) emits signal in both UV and green spectra. **(E):** Cascade Blue dye (i; iv) following transfer. Two additional cells with dye that appear to lack *DsRed* or *eGFP* signals (yellow arrows).

#### EC and PC Intercellular Communication

To determine if PCs and ECs (or precursors) were coupled during vasculogenesis, we used a microinjection dye transfer assay at day 7 of differentiation, when both reporters were detected but before tube formation (Figure 6C-E). We filled *Ng2:DsRed*+ PCs with the intracellular dye Cascade Blue (Figure 6D), which emits in the ultraviolet (UV) range, though it extends into green (e.g. eGFP) wavelengths. We addressed potential green signal overlaps by acquiring images before and after the microinjection (Figure 6C and 6E). As the microinjected PC filled with dye, an adjacent PC began to emit a signal in the UV range, indicating PC-PC coupling (Figure 6D). Subsequent injection of this second PC unfortunately compromised its viability. After allowing 30 minutes for dye transfer, we assessed Cascade Blue fluorescence within this region of interest by confocal microscopy (Figure 6E). Successful dye transfer was detected from the microinjected PC to two adjacent *Flk-1:eGFP*+ cells, consistent with direct communication from the PC to nascent ECs via GJs. We observed dye signal in two additional cells (Figure 6E, i and iv). One of these the cells was in close proximity to a Cascade Blue-*Flk-1:eGFP-*double positive cell, suggesting either transfer directly from the patched PC or via an indirect intercellular route. These data demonstrate direct communication between PCs-ECs and PCs-PCs via GJs prior to vessel tube formation.

### In Early Vascular Development, PCs and ECs Express and Localize Connexin43 at the PC-EC Interface

We assessed FACS-sorted PC (*Ng2:DsRed*+) and EC (*Flk-1:eGFP+)* expression of GJ connexin isoforms in undifferentiated, day 7 and day 10 differentiating DR-ESCs via RNA-Seq (Figure 7A). During differentiation, both cell populations up-regulated *Cx43/Gja1* transcripts to the exclusion of all other mapped connexins. These findings were corroborated in a distinct wild-type (WT) ESC line at day 10 differentiation via 1) cell-specific mRNA expression in ECs and PCs isolated by magnetic cell separation (MACS) (Figure 7B), 2) positive Cx43 labeling at PC-EC interface GJs (Figure 7C), and 3) total protein via Western Blot (Figure 7E). Further, as phosphorylated Cx43 species correspond to differential electrophoretic isoforms (Solan and Lampe, 2018), we examined cell-specific phosphorylation states reflected in Cx43 immuno-blots (Figure 7D). Quantification of the P0-P3 bands in MACS-sorted PCs revealed a distinct increase in the P1 band (Figure 7F) relative to ECs, corresponding to phosphorylation at S365, which indicates localization of Cx43 at the plasma membrane and protection against GJ down-regulation (Solan et al., 2007).

**Figure 7.**
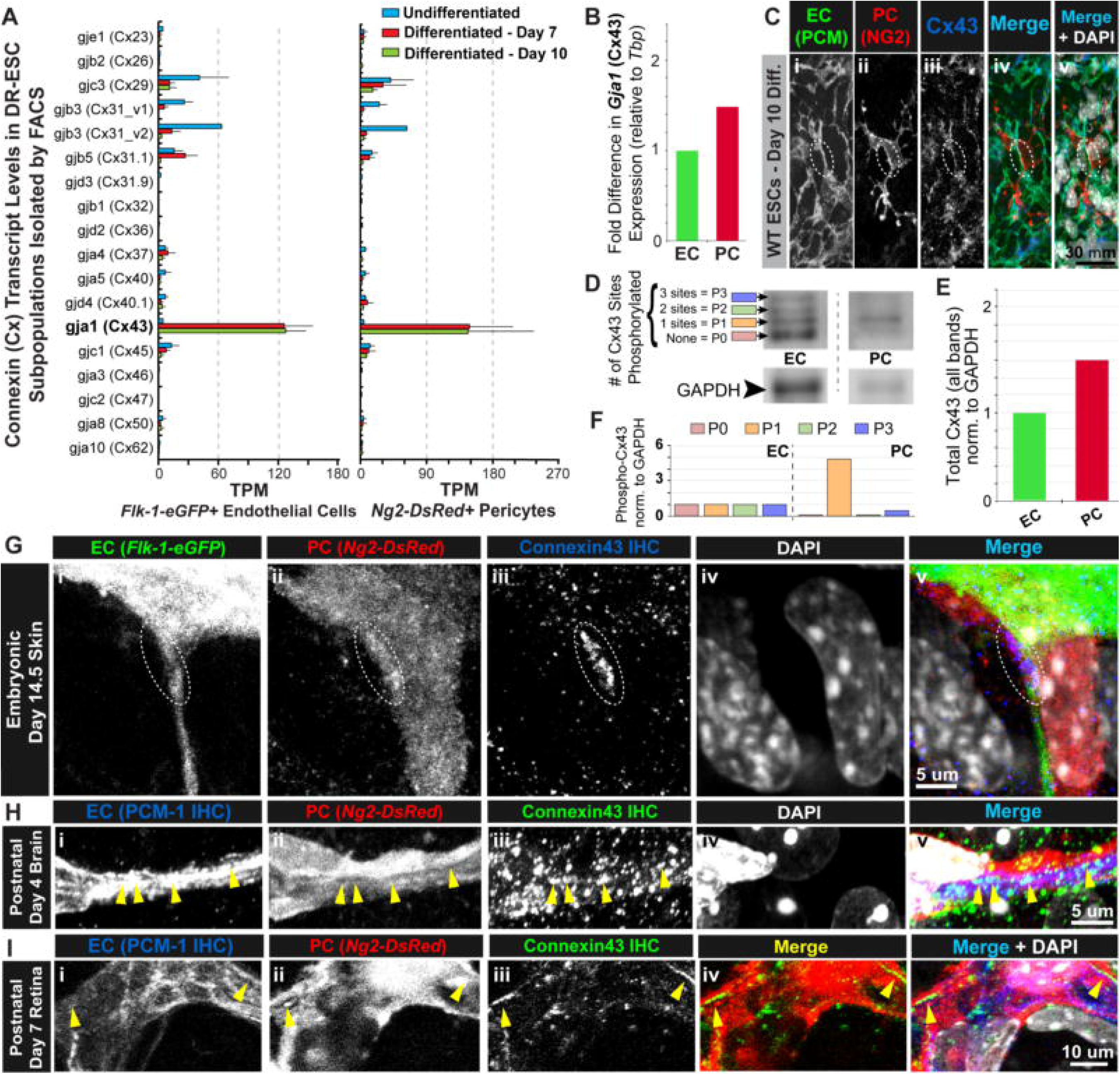
In Early Vascular Development, PCs and ECs Express and Localize Connexin43 at the PC-EC Interface. **(A):** Transcript levels of cell-specific Connexin (Cx) isoforms from FACS-sorted (*Flk-1-eGFP*+ and *Ng2-DsRed*+) subpopulations of undifferentiated, days 7 and 10 differentiated DR-ESCs. TPM, Transcripts per million. Error bars, SD, (n=2). **(B):** Fold difference in *Gja1* (Cx43) in MACS-sorted NG2+ PCs (PCs) vs. PECAM-1/CD31+ ECs (ECs) from WT-ESCs via qRT-PCR. **(C):** Day 10 differentiated WT ESCs labeled for PECAM-1+ (PCM) ECs (i; iv, v), NG2+ PCs (ii; iv, v), and Cx43 (iii; iv, v). Nuclei, DAPI (v). **(D):** Westerm Blot for Cx43 and GAPDH (housekeeping) from PECAM-1/CD31+ ECs and NG2+ PCs isolated from day 10 WT-ESC-derived vessels by MACS. **(E):** Fold difference in total PC Cx43 protein vs. EC Cx43 levels, summing intensities for all Cx43 bands (panel D). **(F):** Fold difference in each Cx43 phospho-isoform from NG2+ PCs vs. CD31+ ECs isolated by MACS. **(G):** Embryonic day 14.5 (E14.5) skin vessels from a *Flk-1-eGFP; Ng2-DsRed* embryo, visualizing ECs (i; v) and PCs (ii; v), and labeled Cx43 (iii; v), which formed GJ plaques at the PC-EC interface (dashed white oval). Nuclei, DAPI (iv, v). *See* ***Figure S7.* (H):** Postnatal day 4 (P4) brain microvessels from an *Ng2-DsRed*+ mouse (PCs, ii; v) labeled for PECAM-1+ ECs (i; v) and Cx43 (iii; v). Nuclei, DAPI (iv; v). Cx43-enriched plaques at the PC-EC border (yellow arrowheads). (I): P7 mouse retina microvessels labeled for EC PECAM-1 (i; v), PC surface NG2 (ii; v), and Cx43 (iii; v). Nuclei, DAPI (iv; v). Cx43 GJ plaques at the PC-EC interface (dashed yellow ovals).

To verify *in vivo* that Cx43 is present at the PC-EC interface, we fluorescently labeled Cx43 in “double reporter” mice at E14.5 in dermal vessels using high-power confocal microscopy (Figure 7G and Figure S7). We observed Cx43 GJs and Cx43- enriched GJ plaques at the interface between *Ng2:DsRed+* PCs and *Flk-1:eGFP+* ECs, bridging across two early-stage vessels. We also examined post-natal day 4 (P4) brain vasculature from an *Ng2:DsRed* mouse. Immunolabeling for PECAM-1 and Cx43 (Figure 7H) revealed numerous Cx43 GJs located at discrete locations along PC-EC borders. We also labeled PECAM-1 and Cx43 in developing P7 *Ng2:DsRed* retina vessels (Figure 7I). At this stage, ECs sprout from a single vessel layer along an astrocyte template (Chappell et al., 2019) with minimal extensions into underlying tissue. We observed Cx43-enriched GJ plaques localized at the PC-EC interface, consistent with our data across stages and tissues.

## DISCUSSION

Pericytes have recently received considerable attention regarding contributions to microvascular function in various tissues and pathologies. While PC are appreciated as critical to stabilization, maturity and permeability of the microvasculature (Armulik et al., 2010; Daneman et al., 2010; von Tell et al., 2006), comprehensive understanding of PC function is far from complete and is predominantly restricted to established or angiogenic vasculature. Here, we used complementary approaches to examine PCs in vasculogenesis. We uncovered fundamental insights into the timing of PC and EC lineage commitment and direct PC-EC communication prior to vessel formation.

We examined EC (*Flk-1:eGFP)* and PC (*Ng2:DsRed*) reporter mouse embryos when vessel structures had not yet begun to form. At embryonic day 8.5 (E8.5) we found distinct *Ng2:DsRed*+ cells among arising *Flk-1:eGFP+* ECs. Imaging at E9.5 showed *Ng2:DsRed*+ cells along the outflow tract of the heart. While NG2 is commonly used as a mural cell marker, expression can occur in oligodendrocyte progenitor cells, though later in development (Trotter et al., 2010), indicating the observed cells as mural cells. It is widely accepted that pericytes are recruited to blood vessels following endothelial tube formation, so we set out to test these findings. To overcome the challenges inherent to *in vivo* interrogations of vasculogenesis, we developed a stem cell line harboring endogenous reporters for PCs (*Ng2:DsRed*) and ECs (*Flk-1:eGFP*) to complement *in vivo* work. These “DR-ESCs” enable live imaging, fluorescent protein labeling, and cell specific-protein and transcript analysis. Consequently, DR-ESCs provide a valuable mammalian model for studying early vascular lineage commitment, EC (Chappell et al., 2016) and PC cell dynamics, augmenting other cell-based approaches (e.g. organoids, human iPSCs, etc.) that model blood vessels with specific focus on PC contributions to vascular development and barrier function (Jamieson et al., 2017; Stebbins et al., 2019).

We validated that DR-ESCs were genetically stable, highly proliferative, and pluripotent. Transcription profiles demonstrated robust EC and PC differentiation within DR-ESCs over time. EC gene expression was consistent with previous observations of ECs arising from angioblasts in mouse ESCs, comparable to developmental processes *in vivo* (Vittet et al., 1996). Interestingly, expression of *Pecam1/CD31* in undifferentiated mouse ESCs, consistent with previous reports (Li et al., 2005; Robson et al., 2001), implicate this cell-cell adhesion molecule in early-development roles prior to EC expression.

We verified that reporters labeled the expected cell populations via antibody labeling and confocal imaging. PCs express *Ng2* across a range of developing vascular beds (Ozerdem et al., 2001), but they are most confidently identified through a combination of labels and morphological features, due to the lack of a singular PC marker (Armulik et al., 2011). We confirmed co-labeling of DsRed+ cells for PC markers, PDGFRb and desmin, and confirmed active NG2 protein synthesis to eliminate the possibility of fluorescence in progeny cells. *Ng2/Cspg4* expression has been described for neuroectoderm-derived glial cells, specifically oligodendrocyte progenitor cells (OPCs), although later in development (Trotter et al., 2010). There was no IHC signal overlap of OPC marker, A2B5, IHC with *Ng2:DsRed*+ cells, although A2B5 fluorescence was observed near to, but not coinciding with, *Flk-1:eGFP+* vessels (Figure 3Bv-xii). These observations indicate that OPCs arise within differentiating DR-ESCs, and suggest the utility of this cell line in studying early stage OPC differentiation alongside the developing vasculature (Maki, 2017). RNA-seq analysis of OPC markers further confirmed PC identity. Interestingly, we observed no co-localization of PECAM-1 or *Flk-1:eGFP+* reporter with NG2-associated signals, although such overlap has been reported for iPSC systems (Kumar et al., 2017), suggesting important distinctions between ESC and iPSC differentiation. Of note, we did not observe double positive cells in E8.5 *Flk-1-eGFP, Ng2:DsRed* embryos. Likewise, as additional cell types such as cardiomyocytes (Ema et al., 2006) and retinal ganglion cells (Okabe et al., 2014) may express Flk-1, we verified *Flk-1-eGFP*+ EC identity via RNA-seq analysis and co-labeling with PECAM-1, and also confirmed that eGFP+ cells were not progeny via Flk-1 ICC throughout differentiation. We confirmed that genetic profiles of differentiating DR-ESCs, by both bulk and FACS-sorted populations, reflected key transcriptional features of vascular development. Together, these quantifications substantiate that: (i) DR-ESC-derived *Ng2:DsRed*+ cells are vascular PCs and/or their precursors, and (ii) these cells engage with nascent endothelium, identified as highly-expressing *Flk-1:eGFP+* cells. These data instill confidence in using these DR-ESCs to study differentiation dynamics and morphological patterns that emerge during the earliest stages of blood vessel formation.

To ensure relevance of DR-ESCs as a model in which to examine vasculogenesis events, we verified that differentiating DR-ESCs give rise to basic vessel structures (Bautch et al., 1996; Doetschman et al., 1985; Marchetti et al., 2002; Yamashita et al., 2000; Yurugi-Kobayashi et al., 2003) within or adjacent to primordial cardiac tissue. This reflects embryonic formation of major blood vessels (Hislop, 2002; Strilic et al., 2009) that occurs through *in situ* coalescence of ECs, thereby establishing conduits connecting the developing heart with the systemic circulation. Long-term, dynamic confocal imaging of differentiating DR-ESCs showed primitive vascular networks that formed *de novo*, quantifiably expanded and ultimately stabilized. Recapitulating *in vivo* events, these DR-ESCs represent an *in vitro* platform for studying the earliest transcriptional and morphological stages of cardiovascular development, which is difficult to achieve within embryonic tissues, and is often limited with respect to real-time imaging approaches.

An advantage of using the *Flk-1* promoter to drive eGFP reporter expression in this DR-ESC line is the ability directly visualize EC differentiation from mesodermal precursors (Pardanaud et al., 1996), which both express *Flk-1* (Ema et al., 2006; Yamaguchi et al., 1993) at differing levels of intensity – an observation mirrored in E8.5 double reporter mouse embryos. Through PECAM-1 labeling we confirmed that high eGFP fluorescence (*Flk-1:eGFP+^high^*) exclusively distinguished ECs from low eGFP fluorescence (*Flk-1:eGFP+^low^*) observed in EC-precursor mesoderm in DR-ESCs. The unique capability to directly observe differentiation dynamics in our model allowed us to identify key features of mesodermal-to-vascular differentiation and associated cellular behaviors during the initial stages of vasculogenesis/angiogenesis. As some studies suggest a common ESC-derived Flk-1+ precursor generates both perivascular and EC lineages (Chen et al., 2016; Yamashita et al., 2000), we characterized EC differentiation relative to PC lineage commitment. We found no evidence of *Ng2:DsRed*+ cells arising from a *Flk-1:eGFP+* cell, and no instances of overlap between *Ng2:DsRed* and *Flk-1:eGFP* fluorescent signals in DR-ESCs or in E8.5 double-reporter embryos, consistent with previous observations (Crisan et al., 2008; Darden et al., 2018; Kim et al., 2016; Zhao et al., 2018). Differences in experimental conditions may explain the discrepancy between studies. Nevertheless, our findings suggest that, while terminally differentiated ECs arise from *Flk-1+* mesoderm, the PC lineage likely emerges from Flk-1-negative precursors, and suggest the need to reconsider reports that PCs arise from an endothelial lineage during development.

PC differentiation and vessel-association are thought to occur subsequent to EC differentiation (Ando et al., 2016; Bergers and Song, 2005; Marmé and Fusenig, 2008; Stratman et al., 2017). Therefore, our initial expectation was that PC differentiation would occur after ECs emerged. Through live-imaging quantification of differentiation dynamics, coupled with transcription profiling, we show that *Ng2:DsRed*+ PCs consistently differentiate prior to *Flk-1:eGFP*+ ECs. These findings suggest that PC differentiation and involvement in blood vessel formation may occur earlier in vascular development than previously appreciated.

During angiogenesis, PCs facilitate remodeling and vessel stabilization (Stratman et al., 2017), and are thought to interact with ECs primarily after early tube and plexus formation occurs (Ando et al., 2016; Bergers and Song, 2005; Marmé and Fusenig, 2008) to allow for greater vascular plasticity (Bergers and Song, 2005). However, live imaging movies of early stage DR-ESC differentiation revealed *Ng2:DsRed*+ PCs that were interacting with *Flk-1:eGFP*+ cells through stellate extensions and cell body contacts. Moreover, *Ng2:DsRed*+ PCs appeared to dynamically engage with surrounding *Flk-1+eGFP*+^high^ ECs, as well las *Flk-1+eGFP*+^low^ mesoderm in a manner suggesting coordination with EC activity, (consistent with observations from PC-HUVEC co-cultures (Zhao et al., 2018)). Dimensional renderings from confocal images of neighboring PCs and ECs supported that they were in direct contact.

To substantiate these findings, we explored whether PCs and ECs were communicating via gap junctions. Gap junctions (GJs), comprised of connexin protein subunits, form at the interface between two cells and often aggregate at higher densities forming plaques, facilitating rapid exchange of small signaling molecules, ions, and 2^nd^ messengers. Cell coupling during the early stages of vascular development has been described previously for mural cells (Hirschi et al., 2003); however, insight into PC-EC GJ communication during vasculogenesis is lacking. At a time point prior to vessel structure formation, we microinjected Cascade Blue into DR-ESC-derived *Ng2:DsRed*+ PCs, which transferred to an adjacent PC and to two adjacent *Flk-1:eGFP*+ cells. These data are supportive of direct PC-EC and PC-PC coupling via GJs during the early stages of vessel development.

Four connexin protein isoforms—Connexin37 (Cx37), Cx40, Cx43, and Cx45— predominate in GJ of mature vasculature (Figueroa and Duling, 2009). Of these, Cx43 is known to be involved in PC differentiation (Hirschi et al., 2003) and vessel maturation (Durham et al., 2015), and plays a key role during epicardial development (Francis et al., 2011; Rhee et al., 2009). During vascular development, mouse Cx43 and Cx45 modulate TGFβ signaling such that disruption of these connexins leads to vascular defects including aberrant mural cell investment (Hirschi et al., 2003; Kruger et al., 2000). RNA-seq analysis of FACS-sorted *Ng2:DsRed*+ PCs and *Flk-1:eGPF*+ EC revealed that *Cx43/Gja1* transcripts were exclusively and prominently upregulated in both cells types throughout differentiation. We confirmed these findings in an additional mouse ESC line (WT-ESCs) at the transcript and protein level, using distinct sorting methods and markers. We examined the functional state of Cx43 (Lampe et al., 2006; Solan and Lampe, 2014) in WT-ESCs, as connexins frequently undergo multi-site phosphorylation that impacts their assembly, function, and stability (Solan and Lampe, 2009). Immunoblotting of sorted PCs favored a migration band that reportedly correlates to 1) increased phosphorylation at S365, 2) primary location at the plasma membrane and a subset of GJs, and 3) protection against down-regulation of GJ channels (Solan et al., 2007). Collectively, these data suggest that Cx43 expression is upregulated in ECs and PCs throughout formation of ESC-derived vessels. Moreover, PC Cx43 appears to favor a phospho-isoform localized to the plasma membrane and limits down-regulation of the channel.

We show presence of Cx43 GJs at the PC-EC interface imaging in E14.5 skin, P4 brain, P7-retina, and differentiating WT-ESCs, via IHC and high-resolution confocal, demonstrating consistency across tissue types and developmental stages. These findings support that PCs and ECs communicate via Cx43-mediated gap junctions, preferentially up-regulating Cx43 during early vessel development. Interestingly, Cx43 knockout mice develop heart outflow tract defects (Huang et al., 2011; Liu et al., 2006; Ya et al., 1998), corresponding to the region of *Ng2:DsRed*+ PC fluorescence along the heart outflow tract of an E9.5 embryo we observed during vasculogenesis.

As current studies focus on PC contribution to numerous pathologies, highlighting their importance in maintaining healthy vessels, these data support a new role for PCs that precedes established or angiogenic vasculature. These findings indicate that PCs arise during vasculogenesis and directly communicate with differentiating ECs prior to vessel structure formation, offering a new fundamental perspective of how blood vessels form. Goals to restore dysfunctional vessels or create new healthy vasculature is focal to a wide range of prominent pathologies such as stroke, Alzheimer’s Disease, **Diabetes** mellitus, heart disease, and cancers. The findings herein provide groundwork for new-phase investigations of PC-mediated vascular function and related pathologies.

## ACKNOWLEDGMENTS

The authors thank the Chappell Lab members for stimulating discussion during manuscript preparation.

## AUTHOR CONTRIBUTIONS

Conceptualization: LBP, JCC; Methodology: LBP, BT, MP, SDP, JCC; Investigation: LBP, BT, LD, SB, AS, JD, HZ, MP, JCC; Writing – Original Draft: LBP, JCC; Writing – Review & Editing: LBP, JCC; Funding Acquisition: KO, HS, JCC; Resources: KO, HS, JCC; Supervision: LBP, JCC

## STAR METHODS

### LEAD CONTACT AND MATERIALS AVAILABILITY

Further information and requests for resources and reagents should be directed to and will be fulfilled by the lead contacts, John C. Chappell, Ph.D. Mouse lines generated in this study have been deposited to the Knockout Mouse Project (KOMP), [pending - name and cat/identifier #].

### EXPERIMENTAL MODEL AND SUBJECT DETAILS

#### In Vivo Animal Models

##### Ethical Statement

All animal work was conducted in accordance with a protocol that was approved by the Virginia Tech Institutional Animal Care and Use Committee (IACUC), which maintains compliance under the Animal Welfare Assurance number A-3208-01 (expiration 07-31-2021).

##### Experimental animals

Will be updated for inclusion, pending revisions.

##### Housing and Husbandry

Mice were housed in a SPF barrier vivarium with individually ventilated caging on Alpha Dri bedding. Maximum density of animals was 5 adults per cage. For our experiments, we housed 2-3 adults per cage, or 1-2 mothers with pups. Animals were on a 12:12 light/dark cycle. Irradiated 2918 and 7904 rodent chow (Envigo) was provided ad libitum. Reverse osmosis water was provided via auto-water system ad libitum. Enviro dry packs and or nestlets enrichment was provided in each cage. The desired room temperature range was 20-22 degrees C with 40-70% humidity. Animal health was assessed daily by dedicated facility animal care technicians. Veterinary care was available 24 hours a day, 7 days a week. Veterinary rounds occurred weekly.

##### Sample Size

Will be updated for inclusion, pending revisions.

##### Allocating Animals to Experimental Groups

Will be updated for inclusion, pending revisions.

#### Cell Lines

##### Wild-type Embryonic Stem Cells (WT-ESCs)

Wild-type Embryonic Stem Cells (WT-ESCs) were a gift from V.L. Bautch (University of North Carolina at Chapel Hill).

##### Double-Reporter Embryonic Stem Cells (DR-ESCs)

*[EC/PC-DR-mESC (RRID:CVCL_XX13)].* DR-ESCs were obtained via blastocyst extraction (see Experimental Procedures in the Method Details section), and therefore no sex was not determined. All cultures were maintained in 5% CO2 at 37°C. DR-ESCs were amplified on irradiated mouse embryonic fibroblasts (iMEFs) in T75-flasks to 80% confluence and cryopreserved in 10% DMSO at passages 8-9. To wean DR-ESCs from iMEFs, cryopreserved cells were thawed and seeded directly onto tissue culture-treated plates without additional iMEFs and passaged several times until iMEFs were visually absent from propagating ESCs. Experiments were subsequently conducted in feeder-free, tissue culture-treated polystyrene without an ECM substrate coating. The DR-ESCs used in this study are pending authentication by ATCC.

##### ESC Maintenance

DR-ESCs were maintained in an undifferentiated state in <u>2i System Medium</u>: Knockout DMEM (without L-glutamine, with sodium pyruvate), 20% heat inactivated (ΔT) FBS, 1% Antibiotic – Pen-Strep, 2mM Glutamine, 1% MEM NEAA, 100 uM β-mercaptoethanol (βME), 1 uM MEK inhibitor, 3 uM GSK inhibitor, 1000 U/mL Esgro LIF (added just prior to using media). Cultures were passaged every 3-5 days, using a 1:3 dilution of 0.25%Trypsin/EDTA in DPBS. WT-ESCs were maintained in an undifferentiated state by using supplementation with medium conditioned by the 5637 human bladder cancer cell line (ATCC #HTB9) that produces LIF, thus eliminating the need for recombinant LIF and the dual-kinase inhibition.

##### ESC Differentiation

To differentiate DR-ESCs, embryoid bodies were gently detached from the dish with 1x dispase and, following 2x washes in DPBS, transferred to <u>Differentiation Medium</u>: high glucose DMEM + gentamycin, 5% FBS, 100x MTG, 1% pen-strep. Differentiation medium was changed every 1-3 days, depending on the day of differentiation and related density of cultures. Differentiation of double reporter ESCs was conducted as described previously (Chappell et al., 2016). WT-ESCs were differentiated as DR-ESCs, with the exception that they were placed in non-adherent plates prior to transfer to tissue-culture treated plates or slide-flasks on day 3 of differentiation.

### METHOD DETAILS

#### Blastocyst Extraction and Stem Cell Isolation

##### Study design

A male *flk-1:eGFP; ng2:DsRed* mouse (*flk-1:eGFP,* Jackson Laboratory #017006 and *ng2:DsRed*, Jackson Laboratory #008241) was bred to a WT female for the purpose of blastocyst harvest and expansion in vitro. A single female was harvested and blastocyst clones were differentiated and screened for *flk-1:eGFP; ng2:DsRed* double-reporter expression.

##### Experimental Procedures

Blastocysts were collected at 3.5 days *post coitus* as previously described (Bryja et al., 2006; Czechanski et al., 2014). After several days of culture on a layer of irradiated mouse embryonic fibroblasts (iMEFs, i.e. non-proliferative feeder cells), individual blastocysts hatched, giving rise to inner cell masses containing pluripotent embryonic stem cells (ESCs). Each colony of ESCs was dissociated and expanded on newly plated feeder cells. Cells were maintained in an undifferentiated state through exposure to LIF, as well as mitogen-activated protein kinase kinase (MEK) and glycogen synthase kinase-3 beta (GSK3b) inhibitors. Colonies were placed in differentiation medium (described above) and, following differentiation, were assessed for *DsRed* and *eGFP* expression using a Zeiss Axio Observer microscope mounted with a Hamamatsu ORCA-Flash 4.0 V2 Digital CMOS camera

##### Experimental outcomes

A single clone was determined to be double-reporter positive. This clone was used for subsequent validation and experimental studies.

#### Embryonic Day 8.5 (E8.5) Harvest and Imaging

##### Study design

A male *flk-1:eGFP; ng2:DsRed* mouse (*flk-1:eGFP,* Jackson Laboratory #017006 and *ng2:DsRed*, Jackson Laboratory #008241) was bred to a WT female in order to harvest embryos at embryonic day 8.5 for confocal imaging.

##### Experimental Procedures

The pregnant dam was euthanized 8.5 days following the plug. The embryos were harvested and placed in glass bottom dishes with DPBS. The embryos were immediately imaged, unfixed, on a Zeiss LSM880 confocal microscope using a 10x or 20x objective.

##### Experimental outcomes

Single plane images were acquired.

#### Karyotyping Analysis

##### Study design

Undifferentiated DR-ESCs were analyzed via karyotyping for chromosomal stability. Twenty-five chromosomal clusters were analyzed from 4 distinct preparations.

##### Experimental Procedures

Undifferentiated double reporter ESCs were arrested in metaphase through exposure to 150ng/mL of Gibco® KaryoMAX® Colcemid^TM^ Solution in PBS (ThermoFisher) for two hours. Cultures were collected, washed, put into single-cell suspension, and treated with 75mM KCl for 20 minutes. Cells were fixed in 3:1 methanol:acetic acid overnight, spun down and re-suspended in fresh fixative twice. Cells were burst by dropping from 2 feet onto angled, cold slides, chilled in 75mM KCL solution, and then briefly heated over steaming water. Slides were dried for 15 minutes at room temperature and DNA of collected cells was labeled with Hoechst 34580 (ThermoFisher). Chromosomes were imaged on a Zeiss LSM880 confocal microscope using a 100x objective, and 2-4 z-axis scans were acquired and compressed, and chromosomes counted.

##### Experimental outcomes

Chromosome numbers were counted from 25 clusters.

#### Alkaline Phophatase Staining

##### Study design

DR-ESCs were analyzed for pluripotency in the undifferentiated and differentiated states, via alkaline phosphatase staining. The experiment was repeated three times, from distinct passages.

##### Experimental Procedures

Undifferentiated DR-ESCs and those differentiated for 5 and 12 days were fixed and stained for alkaline phosphatase (O’Connor et al., 2008) using the StemTAG^TM^ Alkaline Phosphatase Staining Kit (Cell Biolabs). Brightfield images of stained samples were acquired on a Zeiss Axio Observer microscope mounted with a Hamamatsu ORCA-Flash 4.0 V2 Digital CMOS camera

##### Experimental outcomes

Positive vs. negative alkaline phosphatase stains were assessed.

#### Immunocytochemistry

##### Study design

Undifferentiated and differentiated DR-ESCs were assessed for protein markers using immunocytochemistry and confocal microscopy.

##### Experimental Procedures

Cultures were fixed for 15 minutes at room temperature in 4% PFA, and immunostained as previously described (Chappell et al., 2013; Kappas et al., 2008). Primary antibodies (Abs) used were: rat anti-mouse/human Oct3/4 at 1:200, rat anti-mouse platelet-endothelial cell adhesion molecule-1 (PECAM-1)/CD31 (BD Biosciences) at 1:1000, rabbit anti-mouse NG2 (Millipore) at 1:200, and rabbit polyclonal anti-phospho-Histone H3 (PH3, ser10, Millipore) at 1:500, rat anti-mouse PDGFRβ at 1:400, rabbit anti-mouse Desmin at 1:500, mouse anti-mouse A2B5 at 1:100. Secondary antibodies used were donkey anti-rat AlexaFluor647 (IgG; H+L) at ½ the primary Ab dilution factor (Invitrogen), and donkey anti-rabbit AlexaFluor647 (IgG; H+L) at ½ the primary Ab dilution. Incubation with DAPI for 30 min followed all staining. ESC cultures were imaged on a Zeiss LSM880 confocal microscope using a 20x or 40x objective, and 4-12 z-axis scans were acquired and compressed.

##### Experimental outcomes

Immuno-fluorescent signal locations were obtained and visually assessed for spatial relationship to Ng2-DsRed and Flk-1-eGFP fluorescence.

#### RNA isolation, Reverse Transcription and qRT-PCR

##### Study design

Undifferentiated and differentiated DR-ESCs were collected to assess the gene expression levels of endothelial cell and pericyte markers. With the exception of *dsred* and *egfp* expression studies, all experiments were collected from duplicate 60mm tissue-culture treated plates, and biologically replicated 3 times from distinct passages. Cultures harvested for *dsred* and *egfp* studies were harvested from 2 replicates within a single passage. Total RNA was isolated and converted to cDNA, followed qRT-PCR.

##### Experimental Procedures

RNA was isolated using Quick RNA Mini-Prep kit (Zymo). Following purification, 1.5 ug of RNA was reverse-transcribed using High Capacity RT Kit (Invitrogen). Real-time quantitative PCR was conducted in triplicate for each sample, using TaqMan Gene Expression Master Mix (Applied Biosystems) and the Applied Biosystems QuantStudio 6. All targets were normalized to TATA-binding protein (tbp). Primer/probe sets were obtained from Applied Bio-Systems as pre-designed assay mixes, with the exception of the *dsred.T1* primer/probe set that was custom designed through Applied Biosystems. using the following DsRed Express (DsRed.T1) sequence: atggcctcctccgaggacgtcatcaaggagttcatgcgcttcaaggtgcgcatggagggctccgtgaacggccacgagttcgagatcgagggcgagggcgagggccgcccctacgagggcacccagaccgccaagctgaaggtgaccaagggcggccccctgcccttcgcctgggacatcctgtccccccagttccagtacggctccaaggtgtacgtgaagcaccccgccgacatccccgactacaagaagctgtccttccccgagggcttcaagtgggagcgcgtgatgaacttcgaggacggcggcgtggtgaccgtgacccaggactcctccctgcag gacggctccttcatctacaaggtgaagttcatcggcgtgaacttcccctccgacggccccgtaatgcagaagaagactatgggctgggaggcctccaccgagcgcctgtacccccgcgacggcgtgctgaagggcgagatccacaaggccctgaagctgaaggacggcggccactacctggtggagttcaagtccatctacatggccaagaagcccgtgcagctgcccggctactactacgtggactccaagctggacatcacctcccacaacgaggactacaccatcgtggagcagtacgagcgcgccgagggccgccaccacctgttcctgtag

Applied Biosystems assay mixes for the following gene targets are detailed in the Key Resources Table: *Oct4, Fgf4, Sox2*, *Nanog* and *c-kit/CD117, Dab2, Gata6, Actc1* and *Otx2, Pecam1/CD31, VE-Cadherin/Cdh5/CD44, Flk-1/Kdr/Vegfr2, Icam2/CD102, NG2/Cspg4, Pdgfrβ*/*CD140b*, *Desmin*, and *N-cadherin/Cdh2, Tgfβ1*

##### Experimental outcomes

For each biological replicate, the Relative Quantitation (RQ) was calculated from ΔCT values and averaged to determine the reported RQ value. The error was calculated as standard error (where n=3) and standard deviation (where n=2).

#### Flow Cytometry

##### Study design

Undifferentiated and days 5, 8 and 10 differentiated DR-ESCs were processed to assess Ng2:DsRed (pericyte) and Flk-1:eGFP (endothelial cell) reporter overlap via flow cytometry and to demonstrate lack of reporter fluorescence in undifferentiated cells.

##### Experimental Procedures

DR-ESCs in 60mm tissue-culture treated dishes were washed with DPBS and incubated for 30 min at 37°C, 5% CO2 in a mixture of 2.5ug/ml collagenase I and 2.5ug/ml collagenase IV in DPBS. Cells were detached from the dish using a cell scraper and manually dissociated into single-cell suspension in the collagenase mixture. Cells were washed 2x in cold FACS buffer (2% FBS, 1% BSA in DPBS). Cells were re-suspended in cold 2ml FACS buffer and filtered through a 70um filter and then a 40um filter. WT-ESC and single reporter ESCs (Flk-1:eGFP) were grown and harvested alongside DR-ESCs as controls and to establish compensation parameters. Due to lack of an available Ng2:DsRed single reporter ESC line UltraComp eBeads (Invitrogen 01-2222-41) were incubated with PE rat anti-mouse CD31 (BD Pharmingen 553373) to establish a DsRed control, according to manufacturers directions. Flow Cytometry experiments were performed on a Sony SH800 Flow cytometer. FlowJo software was used for analysis.

##### Experimental outcomes

Flow cytometry density plots were created for Ng2:DsRed and Flk-1:eGFP reporter detection in WT-ESCs as compared to DR-ESCs in undifferentiated, day 5, 8, and 10 differentiated cells.

#### Fluorescence activated Cell Sorting (FACS)

##### Study design

Undifferentiated and days 7 and 10 differentiated DR-ESCs were processed and sorted via FACS using endogenous Ng2:DsRed (pericyte) and Flk-1:eGFP (endothelial cell) reporter expression for the purposes of transcriptional analysis via RNA-sequencing. Two biological replicates were obtained from distinct passages.

##### Experimental Procedures

DR-ESCs in 60mm tissue-culture treated dishes were washed with DPBS and incubated for 30 min at 37°C, 5% CO2 in a mixture of 2.5ug/ml collagenase I and 2.5ug/ml collagenase IV in DPBS. Cells were detached from the dish using a cell scraper and manually dissociated into single-cell suspension in the collagenase mixture. Cells were washed 2x in cold FACS buffer (2% FBS, 1% BSA in nuclease free PBS). Cells were re-suspended in cold 2ml FACS buffer and filtered through a 70um filter and then a 40um filter. WT-ESC and single reporter ESCs (Flk-1:eGFP) were grown and harvested alongside DR-ESCs as controls and to establish compensation parameters. Due to lack of an available Ng2:DsRed single reporter ESC line, UltraComp eBeads (Invitrogen 01-2222-41) were incubated with PE rat anti-mouse CD31 (BD Pharmingen 553373) to establish a DsRed control, according to manufacturers directions. FACS was performed on a Sony SH800 Flow cytometer. Following compensation, cells were sorted into nuclease-free PBS and immediately placed on dry ice and subsequently used for RNA-sequencing (Genewiz, NJ).

##### Experimental outcomes

Sorted populations of Flk-1:eGFP+ and Ng2:DsRed+ whole cells were obtained via FACS.

#### RNA-Sequencing

##### Study design

FACS-sorted, whole-cell populations of endothelial cells (Flk-1:eGFP) and pericytes (Ng2:DsRed) obtained from undifferentiated, day 7 and 10 DR-ESC, as well as unsorted populations from the same time points, were input for RNA-sequencing to determine cell-specific gene expression. Two biological replicates were obtained and processed.

##### Experimental Procedures

Whole cells were sent to Genewiz (NJ) for ultra-low input RNA-sequencing via Illumina HiSeq 2×150 bp at approximately 350M paired reads per lane. Raw data was delivered in FASTQ format and aligned to the *mus musculus* genome assembly, GRCm38.p6, and analyzed via Kallisto (Bray et al., 2016). A cutoff of 5 transcripts per million (TPM) was used.

##### Experimental outcomes

Cell-specific (Flk-1:eGFP endothelial cells Ng2:DsRed pericyte) transcript expression was assessed via RNA-sequencing, as transcripts per million (TPM).

#### Short-Acquisition Time-Lapse Imaging

##### Study design

DR-ESCs at day 12 of differentiation were imaged using epi-fluorescence to assess Flk-1:eGFP vessel structures and Ng2:DsRed pericytes relative to contracting cardiomyocytes.

##### Experimental Procedures

Short-acquisition time-lapse imaging (less than 5 mins) was used to image live cultures with a 20x objective on a Zeiss Axio Observer microscope mounted with a Hamamatsu ORCA-Flash 4.0 V2 Digital CMOS camera. Images were acquired at 10 frames per second for 90 seconds.

##### Experimental outcomes

Cultures were visually assessed for presence of vessel structures and pericytes relative to contracting cardiomyocytes.

#### Long-Acquisition Live Confocal Imaging

##### Study design

Ultra-long time-course confocal microscopy was used to obtain real-time videos of differentiating DR-ESCs as vessel structures arise *de novo.* Six separate cultures were used to obtain 21 distinct locations for imaging.

##### Experimental Procedures

Live, long-acquisition imaging (at least 12 hours or longer) of day 6 through 12 differentiating ESCs was conducted on a Zeiss LSM800 confocal microscope (full environmental chamber) using a 20x objective and acquiring images at 23-24 minute intervals. A z-stack of 6-10 images was obtained for each scan with 3-4 microns between focal planes, and each stack was compressed into a single image for each time point. Representative time-lapse sequences shown are from non-consecutive images.

##### Experimental outcomes

Twenty-one confocal live-imaging videos were obtained, spanning days 6-12 of differentiating DR-ESCs.

#### ImageJ/FIJI Vessel Analysis

##### Study design

Vessel structures during early development were characterized in differentiating DR-ESCs (from day 6-12) via analysis of 21 real-time videos obtained from ultra-long time-course confocal microscopy (described above). Movies were analyzed for changes over time in the following parameters: vessel diameter, branch points per mm of vessel length, branch length (mm), and percent of total vessel area. Movies were also analyzed for the timing of emergent *Flk-1:eGFP^low^* mesoderm, *Flk-1:eGFP^high^* endothelial cell and *Ng2:DsRed* pericyte fluorescent signals for the purposes of comparing timing of differentiation of each cell type. Lastly, movies were analyzed for the percent of *Ng2:DsRed* pericytes and *Flk-1:eGFP^high^* endothelial cells which arose from *Flk-1:eGFP^low^* mesoderm precursors. In each case, the researcher conducting the analysis was blinded to the expected outcomes and were not otherwise involved with the study.

##### Experimental Procedures

**Experimental Procedures:** Each movie was analyzed for the parameters described using Image J software. For each parameter, the green channel was isolated and modified into an 8-bit image to render it binary. To find vessel diameter, a line was drawn using the line tool from one side of the vessel to the other at a 90 degree angle to the vessel structure. The length of the drawn line was manually recorded into a spreadsheet. A few vessels were chosen at random for each video and the diameter of these vessels were recorded at multiple time intervals during the video. To find the branch data, the brightness of the image was increased. Using the paintbrush tool, the vessels were traced using black ink to contrast the bright background. The threshold was increased until only the tracing was visible. The image was then skeletonized and analyzed to give branch length and branch points per mm of vessel length. To find percent of total vessel area, the threshold was adjusted to minimize the background signal, and then the area was analyzed using the Measure feature. Measure had been previously set to record the total area and percent area.

##### Experimental outcomes

Branch points per mm of vessel length, branch length (mm), percent of total vessel area, timing of emergent *Flk-1:eGFP^low^* mesoderm, *Flk-1:eGFP^high^* endothelial cell and *Ng2:DsRed* pericyte fluorescent signals, and percent of *Ng2:DsRed* pericytes and *Flk-1:eGFP^high^* endothelial cells that arose from *Flk-1:eGFP^low^* mesoderm precursors, were blindly quantified from live imaging movies of days 6-12 differentiating DR-ESCs.

#### Microinjection Dye Transfer

##### Study design

Direct cell-to-cell communication during early vessel development was assessed via single-cell microinjection of Cascade Blue dye into *Ng2:DsRed+* pericytes in day 8 differentiating DR-ESCs. Two *Ng2:DsRed+* pericytes were microinjected.

##### Experimental Procedures

Undifferentiated DR-ESC embryoid bodies were seeded on 12mm round cover glass and placed in differentiation medium (described above). On day 8 of differentiation, differentiation medium was removed and DR-ESCs were equilibrated in microinjection buffer (1% FBS in DBPS with magnesium and calcium) at 32-33 °C and 5% CO2, approximately 30 minutes prior to the micro-injection experiment. Cells on coverslips were placed onto the perfusion chamber of an upright microscope (Leica DMLFSA) with ×40 water immersion lens. During all the recordings, carbogen-bubbled microinjection buffer was continuously superfused (2 ml/min) to keep cells healthy and avoid any dye accumulation and uptake by damaged cells. All recordings were made at 32–33 °C using an inline feedback heating system (Cat# TC 324B, Warner Instruments). Patch pipettes of 3–5 MΩ open-tip resistance were produced from standard borosilicate capillaries (WPI, 4IN THINWALL Gl 1.5OD/1.12ID) using HEKA PIP 6 vertical pipette puller. Patch pipettes were filled with an intracellular solution of 134 mM potassium gluconate, 1 mM KCl, 10 mM 4-(2-hydroxyethyl)-1- iperazineethanesulfonic acid (HEPES), 2 mM adenosine 5′-triphosphate magnesium salt (Mg-ATP), 0.2 mM guanosine 5′-triphosphate sodium salt (Na-GTP) and 0.5 mM ethylene glycol tetraacetic acid (EGTA) (pH 7.4, 290–295 mOsm). We added 5 µl cascade blue (ThermoFisher #C687, 50 mg/ml stock solution in deionized water) in 1ml intracellular buffer just before the recording. Patch pipettes were visually guided using MM-225 micromanipulator (Sutter Instrument, Navato, CA). We obtained transmission, fluorescence images of individual fluorophore from the selected field of view before and after microinjection of dye, using Axiocam MRm camera attached to the microscope. After making a gega-ohm seal on the patched cell, gentle suction was applied to break the patched membrane and reach into whole cell mode. Whole cell mode was confirmed by checking the cascade blue fluorescence immediately after braking the seal. We allowed cascade blue to diffuse into the cell for 30 minutes and carefully retracted patch pipette to avoid pulling away of the cell with the pipette. Cells were immediately fixed in PBS containing 4% Paraformaldehyde (PFA) (Electron Microscopy Science, cat# 15714-S) for 5 minutes and rinsed 3 times with PBS. Coverslips were mounted (Thermo Fisher, Invitrogen, catalog number: S36936) on glass slides and images were taken immediately at various magnifications using Nikon A1 confocal microscope. The microinjected cells were matched with previously taken fluorescence and transmitted images during the microinjection procedure.

##### Experimental outcomes

Transfer between two *Ng2:DsRed+* pericytes and from an *Ng2:DsRed+* pericyte to and *Flk-1:eGFP+* endothelial cells was observed via confocal microscopy. A second microinjected *Ng2:DsRed+* pericytes did not survive the length of the experiment.

#### Magnet-Assisted Cell Sorting (MACS)

##### Study design

Endothelial cells (PECAM-1+) and pericytes (An2+) were sorted from WT-ESCs via MACs for the purpose of qRT-PCR and immuno-blot analysis of connexin-43 expression. Four to eight biological replicates were collected for qRT-PCR analysis.

##### Experimental Procedures

Pericytes and endothelial cells were enriched from WT ESCs cultures at day 10 differentiation and sorted via MACS (Miltenyi Biotec) as previously described (Chappell et al., 2013; Darden et al., 2018). Briefly, cells were dissociated using dispase and type 1 collagenase (Fisher, Cat #NC9633623) at 37 °C for 40 min, and put into single cell suspension via mechanical pipetting. Cells were washed and filtered through a 70uM filter. Cells were re-suspended in autoMACS® running buffer (MiltenyiBiotec, Cat #130-091-221). Fc-Receptor blocking reagent (Miltenyi Biotech, Cat #130-092-575) was added as per the manufacturer’s instructions. Cells were re-suspended in autoMACS® buffer containing anti-An2 microbeads (MiltenyiBiotec, Cat #130-097-170). An2 is a homologue of Ng2 (Taylor et al., 2010). Labeled cells were passed through QuadroMACS separator LS columns (Miltenyi Biotec, Cat #130-091-051, 30-micron pre-separation filters). Cells flowing through the column but not isolated by the magnetic field were collected for secondary selection of endothelial cells, which were re-suspended in MACS® buffer. Following incubation with Pecam1/CD31 primary antibodies conjugated to R-phycoerythrin (PE) (BD Pharminogen, Cat #553373), cells were then incubated with anti-PE microbeads, as described above. After each MACS column separation, target populations were collected and re-suspended in TRIzol (Invitrogen, Cat #15596018) or RIPA buffer for transcript or protein analysis, respectively. [1X RIPA buffer: 50mM Tris, pH 7.4, 150mM NaCl, 1mM EDTA, 1% Triton X-100, 1% deoxycholate, 0.02mM Na3VO4, 0.01mM NaF, 1x HALT, 10% SDS, 5.55mM NEM].

##### Experimental outcomes

Endothelial cells (PECAM-1+) and pericytes (An2+) were MACS-sorted and collected for transcript and protein analysis.

#### SDS-PAGE and Western Blot

##### Study design

Protein lysates of endothelial cells (PECAM-1+) and pericytes (An2+) obtained by MACS were separated by SDS-PAGE and immuno-blotted using anti-phospho-Cx43 to determine overall protein level and phosphorylation states of Cx43. Connexin 43 expression was normalized to GAPDH.

##### Experimental Procedures

Total protein concentrations were determined using a standard BSA curve (DC Protein Assay, Biorad) according to manufacturer’s instructions. Normalized proteins were separated by SDS-PAGE on 4–15% Mini PROTEAN® Pre-cast TGX gels (Bio-Rad, Cat # 456-1086). Separated proteins were transferred to Immobilon-FL PVDF (EMD Millipore, Cat # IPFL00010) that was subsequently blocked overnight [5% BSA in Tris-buffered solution, 0.1% Tween 20 (TBS-T)] at 4 °C. The membrane was exposed (in TBST) to primary antibodies rabbit anti-mouse phospho-Connexin43 (Cell signaling, Cat # 3511) at 1:1000 and goat anti-GAPDH (Abcam, Cat ab9485) at 1:1000 overnight at 4 °C and to secondary antibodies donkey anti-rabbit AlexaFluor 488 (Jackson ImmunoResearch Cat #711-545-152) and donkey anti-goat AlexaFluor 647 at 1:10,000 (Jackson ImmunoResearch Cat #705-605-147) for 2 h at room temperature. Protein bands were imaged and quantified on a ChemiDoc system (Bio-Rad) using Bio-Rad Image Lab 5.1 software.

##### Experimental outcomes

Connexin43 protein expression and phosphorylation states were quantified for MACS-enriched endothelial cells (PECAM-1+) and pericytes (An2+).

#### Immunohistochemistry

##### Study design

Tissues were collected from mice at embryonic day 14.5 (*Flk-1:eGFP*; *Ng2:DsRed* skin) and post-natal days 4 (*Ng2:DsRed* brain) and 7 (*Ng2:DsRed* retina) and examined for Connexin43 (Cx43) expression patterns via immunolabeling. A single mouse was used for each staining.

##### Experimental Procedures

Postnatal mice were harvested in an isoflurane chamber followed by thoracotomy and, after perfusion with 2% PFA, cervical dislocation. Dissected retina and brain were fixed in 2% PFA overnight and stored at 4°C in PBS. For embryonic skin tissue, the pregnant female was euthanized using standard CO_2_ protocol, the embryos harvested and subject to secondary euthanasia confirmation via decapitation followed by whole body fixation in 4% PFA overnight at 4°C. To prepare tissues for IHC, the p7 retina was processed into leaflets and the p4 was sectioned into xxx um slices using a vibratome. The E14.5 skin section was stained whole. Tissues were blocked and permeabilized for 1h at RT in 1% Triton X-100 in PBS (PBS-T) with 10% Normal Donkey Serum (NDS). Tissues were incubated overnight at 4°C in PBST containing 1:500 rabbit anti-mouse Cx43 and, with the exception of E14.5 skin, also 1:500 goat anti-mouse PECAM-1. The tissues were washed in PBS and incubated for 2h at RT in TBST with 1:500 each of anti-goat AlexaFluor 647 and anti-rabbit AlexaFluor 488. Tissues were washed in PBS. DAPI was added at 1:1000 to the second to last wash and tissues were slide mounted for confocal imaging.

##### Experimental outcomes

Embryonic day 14.5 skin section, postnatal day 4 brain and a postnatal day 7 retina were immunolabeled in the far-red spectrum for Cx43 and, for postnatal tissues, PECAM-1 in the green spectrum to visualize endothelial cells

### QUANTIFICATION AND STATISTICAL ANALYSIS

Statistical details of experiments can be found in the figure legends, including the value of n, what n represents, and dispersion and precision measures. Statistical comparisons were made using GraphPad Prism 6.0, and P-values less than 0.05 were considered significant. Quantitative qRT-PCR RQ values were averaged, and the standard error of the mean (SEM) was found for each group where n ≥ 3 and standard deviation used where n < 3. Each group was compared by one-way ANOVA followed by multiple comparisons via pair-wise Tukey t-test for each gene evaluated

### DATA AND CODE AVAILABILITY

The RNA sequencing datasets generated during this study are available at NCBI Gene Expression Omnibus (GEO) [accession code/web link - pending].

### KEY RESOURCES TABLE

Will be updated for inclusion, pending revisions.

## SUPPLEMENTAL INFORMATION TITLES AND LEGENDS

### SUPPLEMENTAL ONLINE VIDEOS

**Online Video S1.** From *Figure 1B* of the main paper. Time sequence of Ng2-DsRed+ pericytes along the heart outflow tract (indicated in frame 1) of a beating heart in an Ng2-DsRed-positive; Flk-1-eGFP-negative E9.5 mouse embryo. The left and right ventricles are also denoted in frame 1. Scale bar, 200 μm. Time (mm:ss) in upper left corner. Found online at <u>this link</u>.

**Online Video S2.** From the field of view shown in *Figure 4A* of the main paper. Time sequence of “double reporter” (*Flk-1-eGFP* and *NG2-DsRed)* ESC-derived vessels developing within and adjacent to regions of ESCs concurrently differentiating into contractile cardiomyocytes (upper left corner). Time sequence transitions: 1) Brightfield fades to DsRed fluorescence, 2) DsRed fluorescence fades to Brightfield, 3) Brightfield fades to eGFP fluorescence, 4) eGFP fluorescence fades to Brightfield. Scale bar, 300 μm. Time (mm:ss) in upper right corner. Found online at <u>this link</u>.

**Online Video S3.** From *Figure 5A* of the main paper. Single channel (green) time sequence of *Flk-1-eGFP^low^* mesodermal cells giving rise to *Flk-1-eGFP^high^* endothelial cells over days 6-8 of ESC differentiation. Fluorescence intensity levels generated by *Flk-1-eGFP* increase as endothelial cells differentiate from the underlying mesoderm and begin coalesce into basic vessel structures that undergo further remodeling via angiogenic sprouting. Scale bar, 50 μm. Time (hh:mm:ss) in upper right corner. Found online at <u>this link</u>.

**Online Video S4.** From *Figure 5B* of the main paper. Time-lapse confocal images of DR-ESC differentiation over ∼30 hours. EGFP signal intensity in *Flk-1-eGFP+* cells increases over time as endothelial cells (*Flk-1-eGFP^high^*, green) first arise from precursor *Flk-1-eGFP^low^* mesoderm and coalesce into early vessel structures. Fluorescence from *Ng2-DsRed+* pericytes (red) is detected prior to observing signals from *Flk-1-eGFP+* mesodermal and endothelial cells. *Ng2-DsRed+* pericytes appear to interact with the *Flk-1-eGFP+* mesoderm, and these interactions coincide spatially and temporally with endothelial cell organization into primitive vessel-like structures. Scale bar, 50 μm. Time (hh:mm:sec) in upper right. Found online at <u>this link</u>.

**Online Video S5.** From *Figure 6A* of the main paper. Time sequence of *Ng2-DsRed+* pericytes (red) migrating within a region of differentiating *Flk-1-eGFP+* endothelial cells (green) over days 5 to 6 of ESC differentiation. These pericytes appear to engage several endothelial cells during the early stages of their coordination into primitive vascular structures. Scale bar, 50 μm. Time (h:mm) in upper right corner. Found at <u>this link</u>.

**Online Video S6.** From *Figure 6A* of the main paper and *Figure S4* of supplementary material from which representative images were selected. Time sequence of *Ng2-DsRed+* pericytes (red) migrating within a region of differentiating *Flk-1-eGFP+* endothelial cells (green) over days 6 to 7 of ESC differentiation. These pericytes appear to directly engage one another in addition to several nearby endothelial cells during the formation of early-stage vessel structures. Scale bar, 50 μm. Time (hh:mm:ss) in upper right corner. Found online at <u>this link</u>.

**Online Video S7.** From *Figure 6B* of the main paper. Time sequence of an *Ng2-DsRed+* pericyte (red) engaging a *Flk-1-eGFP*+ sprouting endothelial cell (green) during angiogenic remodeling of ESC-derived vessels from days 12 to 13 of differentiation. This pericyte divides, yielding one daughter cell remaining in the original position, and the other daughter cell migrating along the sprout to the newly formed branch point. Scale bar, 20 μm. Time (h:mm) in upper right corner. Found online at <u>this link</u>.

### SUPPLEMENTAL FIGURES

**Figure S1. A Genetically Stable and Highly Proliferative EC- and PC-Reporter ESC Line Maintains Pluripotency, Differentiating into Each Germ Layer. (A):** Karyotyping of undifferentiated mouse DR-ESCs stained by Hoescht. **(B):** Alkaline phosphatase staining of undifferentiated (i), days 5 (ii) and 12 (iii) differentiated DR-ESCs. Scale bar, 100 μm. **(C):** Undifferentiated (i-iii) and day 5 (iv-vi) DR-ESCs labeled for mitotic cells (PH3, green) and nuclei (DAPI, blue). Scale bar, 50 μm. **(D):** Decreased Oct4 expression (red) in DR-ESCs during progression from pluripotency (i-iii) to 5 days (iv-vi) and 12 (vii-ix) of differentiation. Nuclei, DAPI (blue). Scale bars, 50 μm (iii) and 100 μm (vi and ix). **(E):** Stem cell marker expression from undifferentiated (blue), days 5 (red) and 12 (green) differentiated DR-ESCs. **(F):** Germ layer marker expression from undifferentiated (blue), days 5 (red) and 12 (green) differentiated DR-ESCs. *P≤0.05, **P≤0.01, ***P≤0.001, ****P≤0.0001. Error bars, SEM (n=3). *Additional germ layer marker expression by RNA-Seq, Figure S2*.

**Supplemental Figure S2.** From *Figure 1F* of the main paper. **RNA-Sequencing Analysis of Germ Layer Marker Expression in DR-ESCs During Differentiation.** Transcript expression was quantified for additional mesodermal markers (*Brachyury, Bmp4, Foxf1, Snai1*, and *Snai2*), endodermal markers (*Gata4, Sox7,* and *Sox17*) and ectodermal markers (*Fgf8, Foxj3, Nog,* and *Sox1*), in undifferentiated DR-ESCs (blue bars), and days 7 (red bars) and 10 (green bars) of DR-ESC differentiation. Total RNA was generated from unsorted cultures of DR-ESCs, and transcript expression was acquired via paired-end RNA-sequencing. TPM, Transcripts per million. *P≤0.05, **P≤0.01. Error bars represent standard deviation (n=2).

**Supplemental Figure S3.** From *Figure 2D* of the main paper. **Flow Cytometry and Imaging Data Confirming Cell Populations Following FACS. (A):** Flow Cytometry reporter analysis of: (i) undifferentiated DR-ESCs, (ii) day 8 differentiated WT-ESCs (wild type ESCs lacking reporter expression but phenotypically comparable to the DR-ESCs), (iii) day 10 differentiated WT-ESCs, (iv) day 5 differentiated DR-ESCs, (v) day 8 differentiated DR-ESCs, and (vi) day 10 differentiated DR-ESCs. For each density plot (i – vi), *Ng2-DsRed+* event detection is represented on the y-axis and *Flk-1-eGFP+* event detection is represented on the x-axis. Thus, Q1 (quadrant 1, top left) indicates *Ng2-DsRed+* only events, Q2 (quadrant 2, top right) indicates double positive events (i.e. *Ng2-DsRed+* and *Flk-1-eGFP+*), Q3 (quadrant 3, bottom right) indicates *Flk-1-eGFP+* only events, and Q4 (quadrant 4, bottom left) indicates double negative events. The percentage of events that falls inside the quadrant is indicated within each quadrant directly below quadrant name. Total events are shown at the top of each plot. **(B)**: Epi-fluorescent images of day 7 differentiated DR-ESCs acquired prior to (i-iii) and immediately following FACS isolation of *Ng2-DsRed+* (vii-ix) and *Flk-1-eGFP+* (iv-vi) cell populations. Brightfield images (i, iv, and vii) allow for visual comparison of: ***1)*** all cells relative to *Ng2-DsRed*+ cells within the sorted *Ng2-DsRed* (viii) and *Flk-1-eGFP* (v) populations, and ***2)*** all cells relative to *Flk-1-eGFP+* cells within the sorted *Ng2-DsRed* (ix) and *Flk-1-eGFP* (vi) populations. Image magnification for all images, 20x.

**Supplemental Figure S4.** From *Figure 3* of the main paper. **Line Scan Analysis of Endothelial Cell and Pericyte Markers to Establish Cell Identities.** For all panels (A-D), confocal image line-scan analysis (ImageJ/FIJI software) was performed to assess potential overlap of endothelial cell and pericyte IHC marker fluorescence signal with respective *Flk-1-eGFP* or *Ng2-DsRed* reporter signals. Representative regions of signal intensities were quantified by line scans (normalized to background levels for each signal), overlaying cells of interest. Each dotted white line extends from a start position (denoted with an “s”) to an end position (denoted with an “e”) in i, iv and vi of A-D. Normalized fluorescence intensity values are plotted as line graphs (y-axis: Normalized signal intensity and x-axis: distance in µm from “s” to “e”) in iii, vi and ix of A-D. **(A):** From *Figure 3A* of the main paper (merged images iv, viii and xii). Confocal images of days 6-8 differentiated DR-ESC-derived *Ng2-DsRed+* pericytes (i, iv and vii, red) positively co-labeled for pericyte markers PDGFRβ (i-iii, blue), Desmin (iv-vi, blue), and NG2 Immunohistochemistry (NG2-IHC) label (vii-ix, blue). *Flk-1-eGFP+* endothelial cells (vii) are green. DAPI nuclear stain (i and iv) is white. Line scan analysis demonstrates signal overlap of *Ng2-DsRed+* (red lines in ii/iii, v/vi, and viii/ix) with PDGFRβ IHC (blue lines in ii-iii), Desmin (blue lines in v-vi), and NG2 (blue lines in viii-ix). Scale bars, 20 μm (i and iv) and 25 μm (vii). Signal intensities of *Flk-1-eGFP+* signal (green line in ix) and DAPI nuclear stain (white lines in iii and vi) are also represented. **(B):** From *Figure 3B* (merged images iv, viii and xii) of the main paper. Confocal images of DR-ESC-derived *Ng2-DsRed+* pericytes (i, iv and vii, red) void of labeling by the oligodendrocyte precursor cell (OPC) marker A2B5 (i, iv and vii, blue) in day 6 (i), day 8 (iv) and day 10 (vii) differentiated DR-ESCs. *Flk-1-eGFP+* mesoderm and endothelial cells (i, iv and vii) are green. DAPI nuclear stain (i, iv and vii) is white. Line scan analysis demonstrates absence of direct signal overlap between *Ng2-DsRed+* (red lines in ii/iii, v/vi, and viii/ix) and A2B5 IHC (blue lines in ii/iii, v/vi, and viii/ix) across various stages of differentiation. Scale bars, 50 μm (i) and 20 μm (iv and vii). Signal intensities of *Flk-1-eGFP+* signal (green line in ix) and DAPI nuclear stain (white lines in iii and vi) are also represented. **(C):** From *Figure 3C* (merged images iii, vi and ix) of the main paper. Confocal images of DR-ESC-derived *Flk-1-eGFP+* endothelial cells (i, iv, and vii, green) were positively immunolabeled for the endothelial marker PECAM-1 (i, iv, and vii, blue), from day 6 (i), day 8 (iv) and day 12 (vii) of differentiation. Line scan analysis demonstrates signal overlap of *Flk-1-eGFP* (green lines in in ii/iii, v/vi, and viii/ix) and PECAM-1 IHC (blue lines in ii/iii, v/vi, and viii/ix). Scale bars, 20 μm (i and iv) and 50 μm (vii). **(D):** From *Figure 3D* of the main paper. Confocal images of DR-ESC-derived *Flk-1-eGFP+* endothelial cells (i, iv and vii, green) were positively immunolabeled for Flk-1 protein from day 6 (i), day 8 (iv) and day 10 (vii) of differentiation. Line scan analysis demonstrates signal overlap of *Flk-1-eGFP* (green lines in in ii/iii, v/vi, and viii/ix) and Flk-1 IHC (blue lines in ii/iii, v/vi, and viii/ix). Scale bars, 50 μm (i, iv, and iv).

**Supplemental Figure S5.** From *Figure 6A* of the main paper. **Time-Lapse Confocal Images of Pericyte-Endothelial Cell Interactions During Vasculogenic Remodeling**. Representative time-lapse confocal images of *Flk-1-eGFP*+ mesoderm (*Flk-1-eGFP^low^*, ii, v, viii, xi and xiv; green^low^ in iii, vi, ix, xii, and xv) and coalescing endothelial cells (*Flk-1-eGFP^high^*, ii, v, viii, xi and xiv; green^high^ in vi, ix, xii, and xv) over days 6 to 7 of DR-ESC differentiation. *Ng2-DsRed+* pericytes (i, iv, vii, x, and xii; red in iii, vi, ix, xii, and xv) are present before basic organization of endothelial cells, and they appear to engage each other (yellow arrowhead in iv and vi) and also dynamically engage with multiple endothelial cells through ***1)*** stellate extensions (white arrowheads in v-xii) and ***2)*** soma-to-soma (cyan arrows in iv-ix and xiii-xv), as primitive vessel-like structures develop. Scale bar, 50 μm. Time (hh:mm:ss) in upper right. *See **<u>Online</u>*** ***<u>Video S6</u>*** *for full movie from which non-consecutive stills were selected*.

**Supplemental Figure S6.** From *Figure 6B* of the main paper. **Pericyte and Endothelial Cell Engagement During Vasculogenesis in Day 6 and 8 Differentiated DR-ESCs. (A):** Confocal images of individual *Ng2-DsRed*+ pericytes (i, red in iii) in close apposition to individual *Flk-1-eGFP*+ endothelial cells (ii, green in iii) in early stage vasculogenesis. Three distinct *Ng2-DsRed+* pericytes are indicated by solid and dashed cyan arrows and a cyan arrowhead (i and iii). Two distinct *Flk-1-eGFP*+ endothelial cells, engaging with the *Ng2-DsRed+* pericytes, are indicated by solid and dashed magenta arrows (ii and iii). Scale bar, 50 μm. **(B):** Three-dimensional perspectives of z-stacks, shown compressed in ii (and in panel **(A)**), as viewed from the bottom right angle (i) and bottom left angle (iii). Relative intensities of the *Ng2-DsRed* (red) and *Flk-1-eGFP* (green) signals are represented three-dimensionally in an X-Y intensity plot (iv). The three distinct locations of *Ng2-DsRed+* pericytes (correlating between panels **(A)** and **(B)**) are indicated by cyan solid and dashed arrows and a cyan arrowhead (ii and iv). Two individual *Flk-1-eGFP+* endothelial cells, engaging with the *Ng2-DsRed+* pericytes, are indicated by solid and dashed magenta arrows (ii and iv). **(C):** Confocal image of *Ng2-DsRed*+ pericytes (i, red in iii) in close apposition to *Flk-1-eGFP*+ endothelial cells (ii, green in iii), prior to vascular tube formation. Three distinct locations of *Ng2-DsRed+* pericyte engagement with *Flk-1-eGFP+* endothelial cells are indicated by solid and dashed cyan arrows and a cyan arrowhead (iii). Scale bar, 50 μm. **(D):** Orthogonal, single-plane analysis of each of the three distinct locations indicated in **(C)** of *Ng2-DsRed+* pericyte engagement with *Flk-1-eGFP+* endothelial cells (cyan arrowhead in i, cyan arrow in ii, and dashed cyan arrow in iii). Dashed white circles indicate the interactions targeted by the white crosshair lines of the orthogonal single-plane analysis. Orthogonal perspectives of both the vertical and horizontal white crosshair lines are shown at the top and right side, respectively, of each panel (dashed white ovals). **(E):** Three-dimensional renderings of z-stacks, shown compressed in ii (and in panel **(C)**), as viewed from the bottom right angle (i) and bottom left angle (iii). The intensities of the *Ng2-DsRed* (red) and *Flk-1-eGFP* (green) signals are represented three-dimensionally in an X-Y intensity plot (iv). The three distinct locations of *Ng2-DsRed+* pericyte engagement with *Flk-1-eGFP+* endothelial cells (correlating between panels **(C)**, **(D)**, and **(E)**) are indicated by cyan solid and dashed arrows and a cyan arrowhead (ii and iv).

**Supplemental Figure S7.** From *Figure 7G* of the main paper. **Dimensional Z-stack Analysis of Connexin43 at the Endothelial Cell-Pericyte Interface In Embryonic Tissue. (A):** Representative confocal image of embryonic day 14.5 (E14.5) skin capillaries from a *Flk-1-eGFP; Ng2-DsRed* embryo and immunolabeled for Connexin43 (Cx43), facilitating visualization of endothelial cells (green), pericytes (red), and Cx43 (blue). Z-stack of confocal images are shown as compressed and two-dimensional. Scale bar, 5 μm. **(B):** Three-dimensional (3D), angled rendering of a confocal imaging stack, with a higher magnification view of the pericyte-endothelial cell interface in **(C)**, and a further magnified view in **(D)**. Staining of Cx43 revealed a gap junction plaque at the pericyte-endothelial interface (dashed black oval) as well as non-plaque associated Cx43 interspersed along the pericyte-endothelial interface (orange arrowheads in **(C)** and **(D)**). From angled, 3D analysis of the entire z-stack **(B)**, two targeted cross-sections were selected for enhanced resolution imaging – higher magnification (**(B)** and **(C)**, dashed cyan rectangle) and additional magnification (in **(B)**, **(C)**, and **(D)**, dashed magenta rectangle).

