## Supplementary figures and images for "Pericytes Directly Communicate with Emerging Endothelial Cells During Vasculogenesis"

### Figure S1

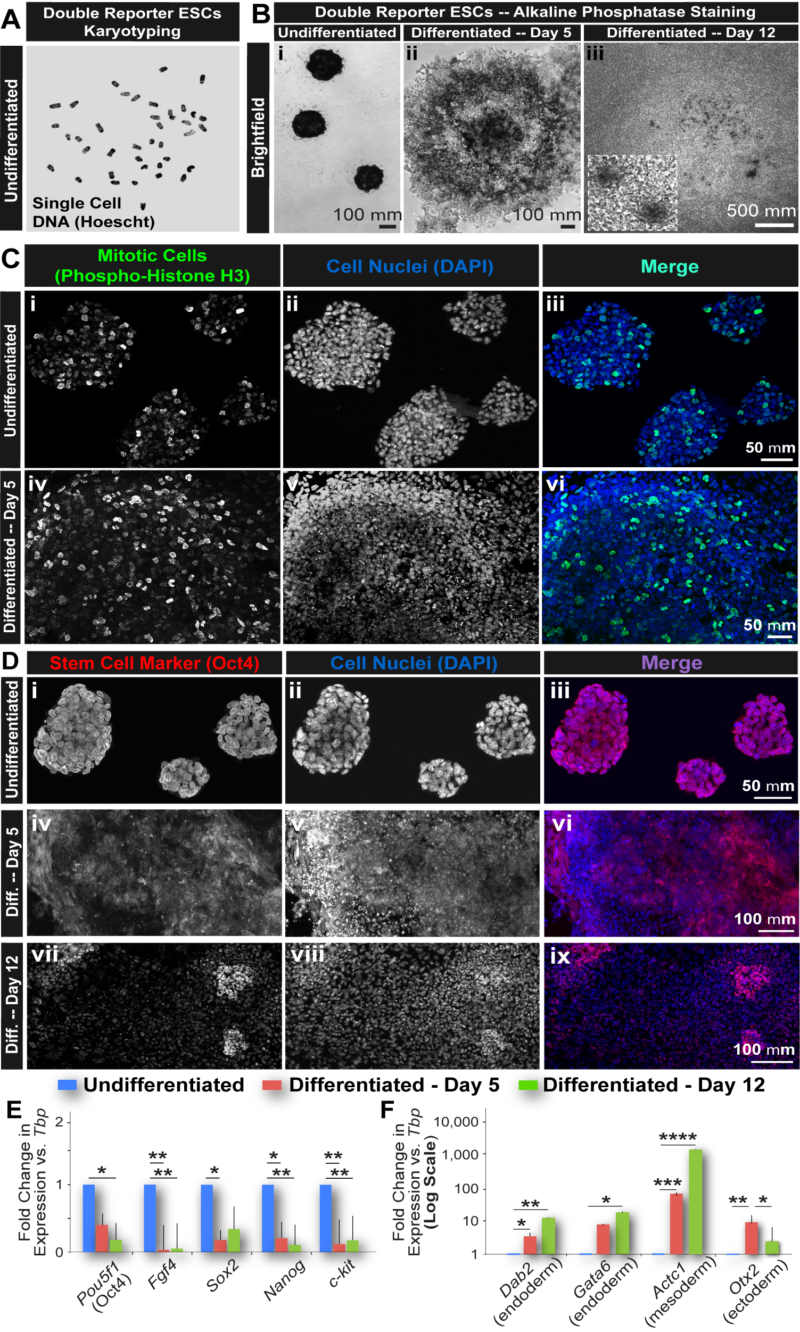

### Figure S2

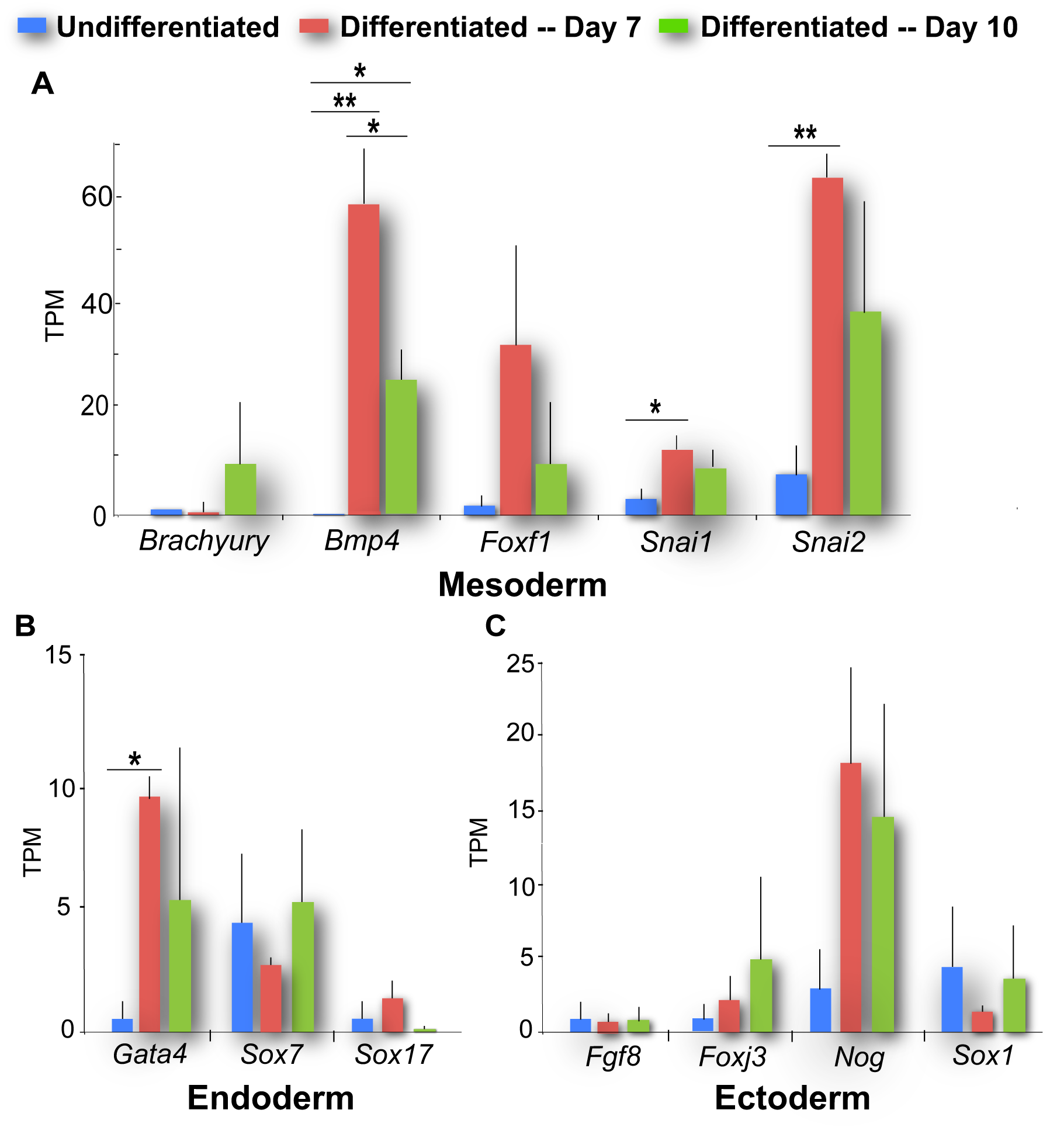

### Figure S3

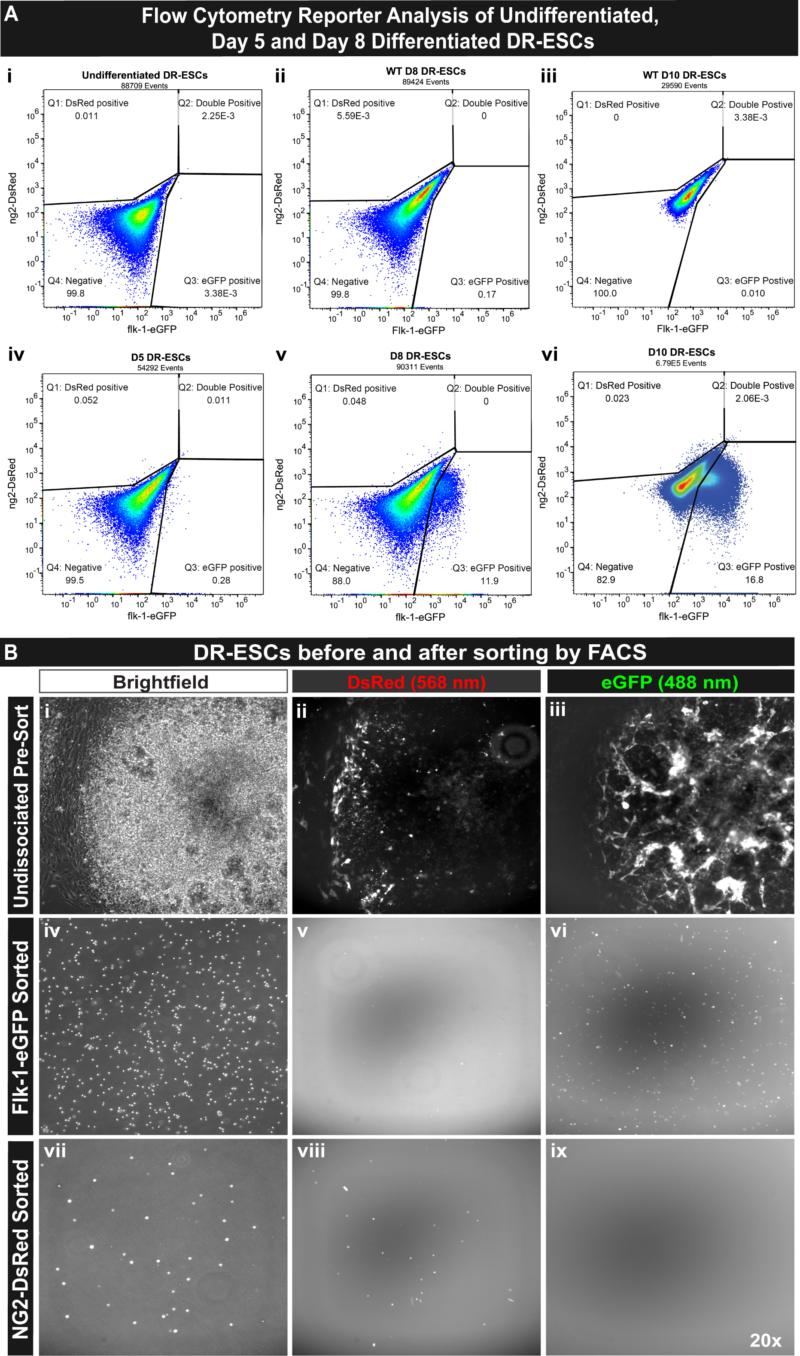

### Figure S4

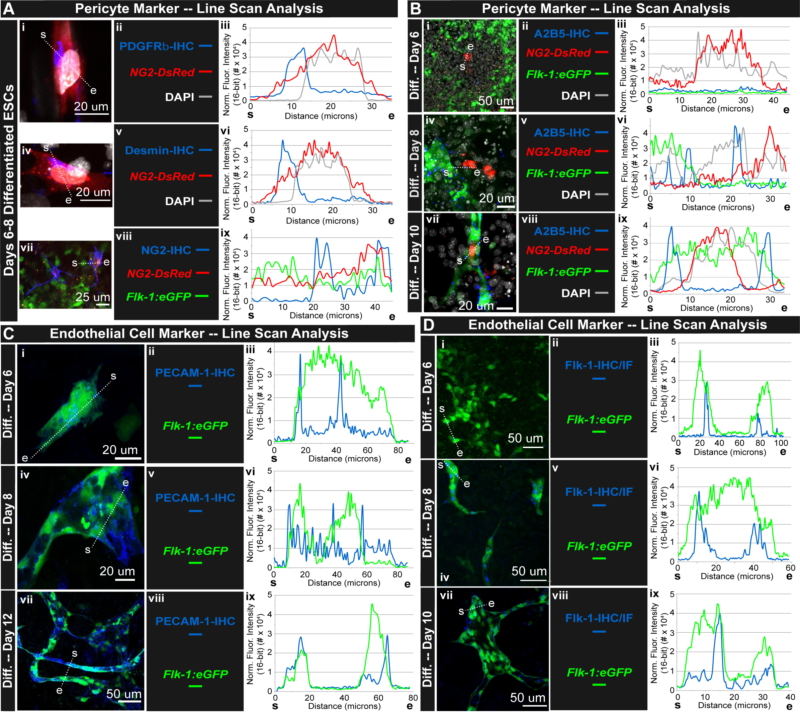

### Figure S5

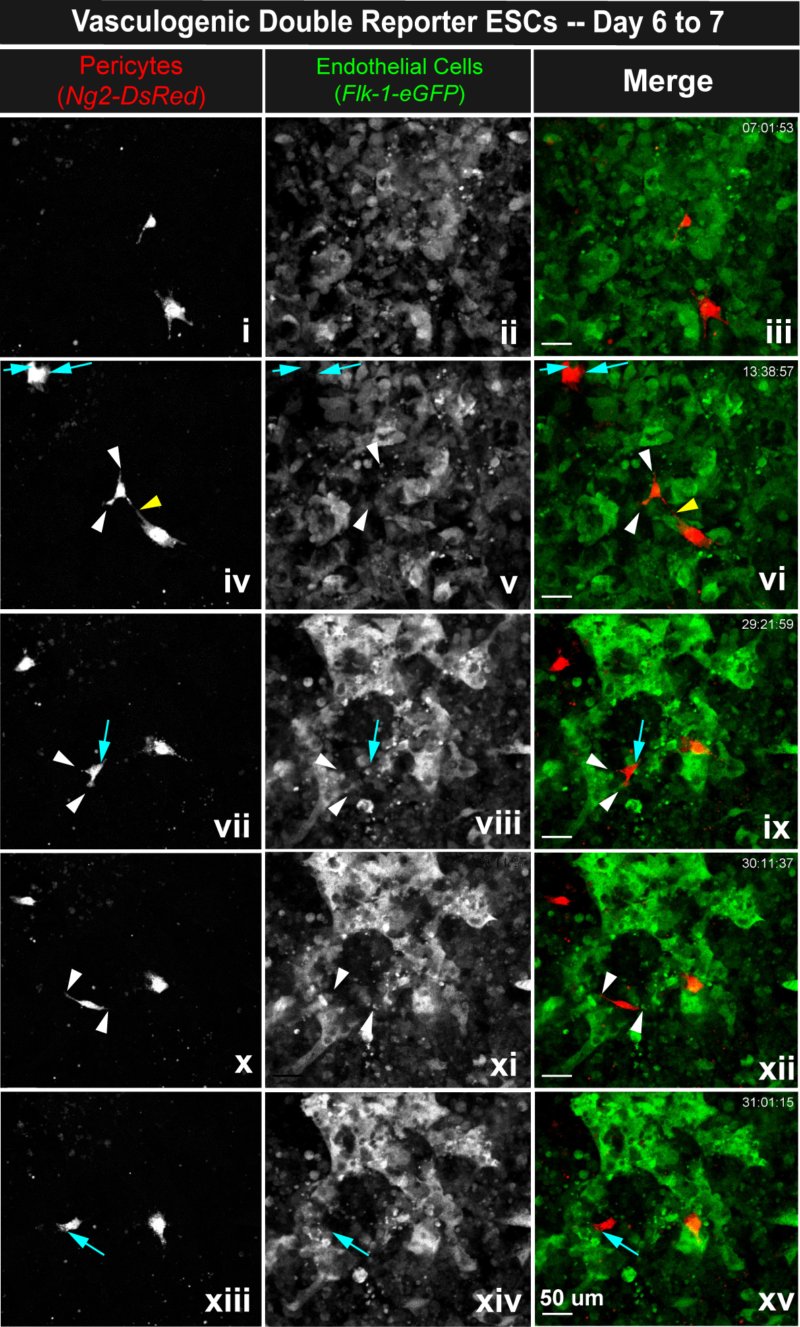

### Figure S6

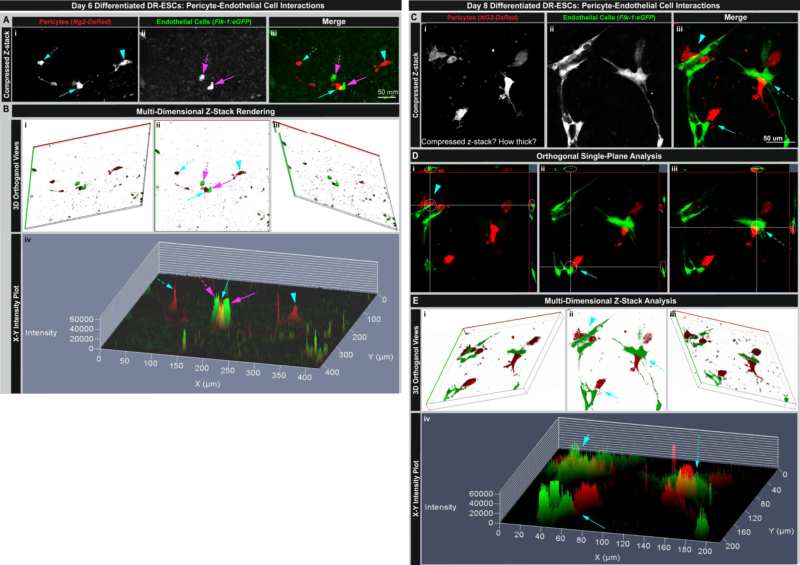

### Figure S7

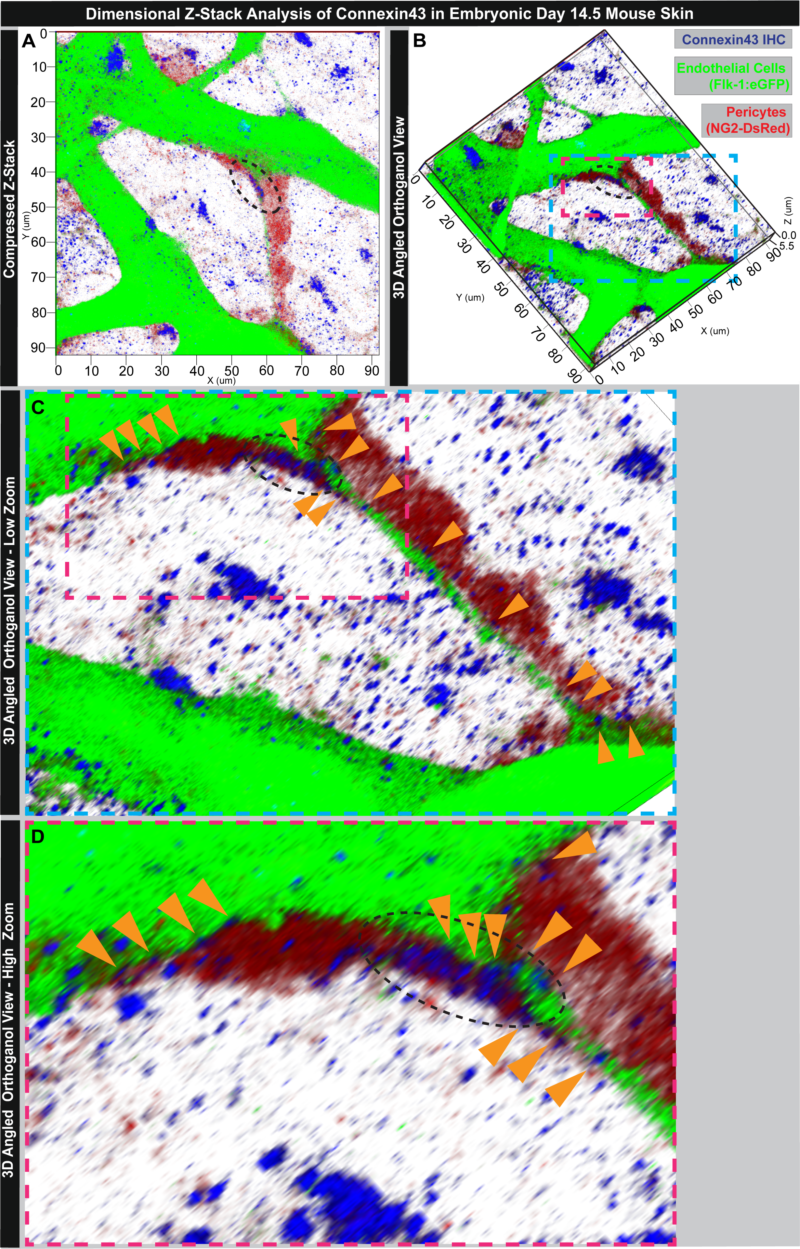
